# Parallel streams define the temporal dynamics of speech processing across human auditory cortex

**DOI:** 10.1101/097485

**Authors:** Liberty S. Hamilton, Erik Edwards, Edward F. Chang

## Abstract

To derive meaning from speech, we must extract multiple dimensions of concurrent information from incoming speech signals, including phonetic and prosodic cues. Equally important is the detection of acoustic cues that give structure and context to the information we hear, such as sentence boundaries. How the brain organizes this information processing is unknown. Here, using data-driven computational methods on an extensive set of high-density intracranial recordings, we reveal a large-scale partitioning of the entire human speech cortex into two spatially distinct regions that detect important cues for parsing natural speech. These caudal (Zone 1) and rostral (Zone 2) regions work in parallel to detect onsets and prosodic information, respectively, within naturally spoken sentences. In contrast, local processing within each region supports phonetic feature encoding. These findings demonstrate a fundamental organizational property of the human auditory cortex that has been previously unrecognized.

## 1 Introduction

A fundamental goal in the neurobiology of language is to understand how acoustic information in speech is transformed into meaningful linguistic content. Speech is thought to be serially processed through the hierarchical structure of the auditory system, from acoustic to phonemic to word and higher order representations (Bizley and Cohen, 2013; DeWitt and Rauschecker, 2012; Leaver and Rauschecker, 2010; Rauschecker and Scott, 2009; Wessinger et al., 2001). As a result, most traditional approaches have been model-based, usually examining the relationship between a well-defined stimulus feature and neural activity. For example, the basic cochlear decomposition of different sound frequencies is reflected in the tonotopically organized maps found throughout the majority of the ascending auditory system, including the primary auditory cortex and adjacent areas (Da Costa et al., 2013; Kaas and Hackett, 2000; Moerel et al., 2012; Saenz and Langers, 2014). In contrast, in the higher order auditory cortex, including the superior temporal gyrus (STG), there is evidence for the encoding of acoustic-phonetic features (Howard et al., 2000; Hullett et al., 2016; Mesgarani et al., 2014; Schönwiesner and Zatorre, 2009). While productive, these approaches often require *a priori* knowledge of potential acoustic (e.g. spectrotemporal) or linguistic (e.g. phoneme, syllable) features.

A major limitation of such model-based approaches is that we do not yet fully know all the potential stimulus features. Debates persist in the linguistics literature regarding the role of phonemes, syllables, and other theorized cognitive representations in the neural processing of speech (Hickok, 2014; Nearey, 2001; Sussman, 1984). Predicting neural responses from reduced sets of features represents a major challenge for characterizing high-order sensory cortices, where neural responses are driven more strongly by complex natural stimuli than their component features. Indeed, recent evidence suggests spectrotemporal modulation tuning to speech in the human STG, yet non-speech control stimuli designed specifically to probe modulation features did not drive strong responses (Hullett et al., 2016; Schönwiesner and Zatorre, 2009).

For these reasons, we used an unbiased, data-driven approach to discover the major patterns of variability in auditory cortex to natural continuous speech. This alternative, model-independent strategy allowed us to identify functional response types across participants, without imposing assumptions about which features or dimensions of speech are most relevant, or about even their localization. To this end, we examined a massive dataset of direct cortical recordings from human participants who were implanted with high-density intracranial electrodes for surgical treatment of epilepsy. Participants listened to natural speech sentences while recordings were made from the entire auditory cortical pathway, including core, belt, and parabelt auditory cortex, as well as related prefrontal and motor areas known to be involved in speech perception.

We first applied unsupervised non-negative matrix factorization (NMF) to the neural response profiles of 2,100 electrodes across 27 participants listening to natural sentences. We discovered two canonical response profiles that divided speech-responsive cortex into spatially distinct processing zones: a caudal zone dominated by strong responsivity to stimulus onsets, and a rostral zone that was activated in parallel and showed generally sustained activity throughout the stimulus. These different response profiles observed in STG electrodes were also observed throughout the entire speech-responsive cortex, including early auditory cortical areas on the temporal plane as well as prefrontal and motor areas. Segmental phonetic features were represented locally at single electrodes, and were embedded equally in each zone. Together, these parallel streams define a striking pattern of temporal dynamics that govern the auditory processing of speech.

## 2 Results

### 2.1 Human superior temporal gyrus is partitioned into two zones with distinct sentence-level response profiles

Participants listened to 499 naturally spoken sentences from the TIMIT acoustic-phonetic corpus, spoken by 402 male and female talkers. We applied an unsupervised soft clustering algorithm, convex non-negative matrix factorization (cNMF), on recordings from 2,100 electrodes across these patients, using the high gamma time series from all speech-responsive electrodes throughout the recording session (see Methods). This analysis was designed to define the electrode response profiles that were similar across patients, and did not rely on the identification of any acoustic or phonetic segmentation, nor knowledge of spatial location or anatomical area of the recordings. Our analysis showed that two dominant response profiles characterized the activity of the electrodes

To understand the differences between these response types, we began by visually inspecting the response profiles at individual electrodes to single sentences (Fig 1a, also see Supplemental Fig. 1). We observed a striking difference in responses: one group showed very strong responses to sentence onset, and the other appeared to have responses that were sustained or had broad peaks at various times throughout the sentence. We then examined the response across the entire population, by plotting the cluster-weighted average responses to single sentences, aligned by sentence onset and sorted by length (Fig. 1b). Because the cluster-weighted time series is collapsed across all electrodes within a cluster, only the overall shape of population activity is observed. This “onset” and “sustained”-like response profile is thus a general characterization of the two populations of electrodes.

Although our unsupervised analysis was designed to uncover similarities in functional structure across brain areas and across subjects, we also wanted to examine if these functional properties were spatially localized. The first cluster that was sensitive to onsets was mainly localized to posterior STG, whereas the other was localized to the anterior and middle STG (Fig 1c). This spatial segregation was not a requirement of our clustering algorithm, which was performed on all subjects simultaneously without *a priori* information about electrode locations. Spatial organization of response types was clearly seen in both left (*N* = 17 subjects) and right (*N* = 10 subjects) auditory cortices across all participants (Fig. 1d, also see Supplemental Fig. 2). We henceforth refer to these as Zone 1 and 2 by analogy to a distinction in the animal brainstem literature (Cant and Benson, 2007), which may be the origin of the ascending connectivity to these cortical zones. In addition to the large number of electrodes with sustained-type responses in Zone 2, we also observed a limited number of sustained-type electrodes posterior to Zone 1, however this appeared to be highly variable across individuals.

We quantified functional and anatomical clustering strength using the silhouette index, which measures the degree of within-cluster and between-cluster similarity. A silhouette index close to 1 indicates good clustering. The degree of both functional and anatomical clustering within Zone 1 and 2 was significantly higher than chance, providing evidence that electrodes belonging to each group were distinct processing zones (Fig. 1e, *p* < 0.001, Wilcoxon signed rank tests).

These clusters represent the major source of variance within our dataset. Similar clustering was observed regardless of clustering method (for example, K-means and other factor analytic methods showed similar results; data not shown). Across all subjects, the two clusters explained 14.5% of the variance in the data. Adding more clusters explained only marginally more variance (Supplemental Fig. 3a). More importantly, within the additional clusters we observed the same “onset” and “sustained” response types, mostly further subdivided according to response magnitude (Supplemental Fig. 3b-d).

At a global level, these results suggest that distinct regions of the human STG are sensitive to important temporal cues in sentences, such as onset and ongoing speech. However, we also know from previous work that STG is sensitive to spectrotemporal and phoneme feature cues in speech (Hullett et al., 2016; Mesgarani et al., 2014). To relate these findings, we next wanted to know how processing in each zone related to other acoustic and phonetic feature representations.

**Figure 1:**
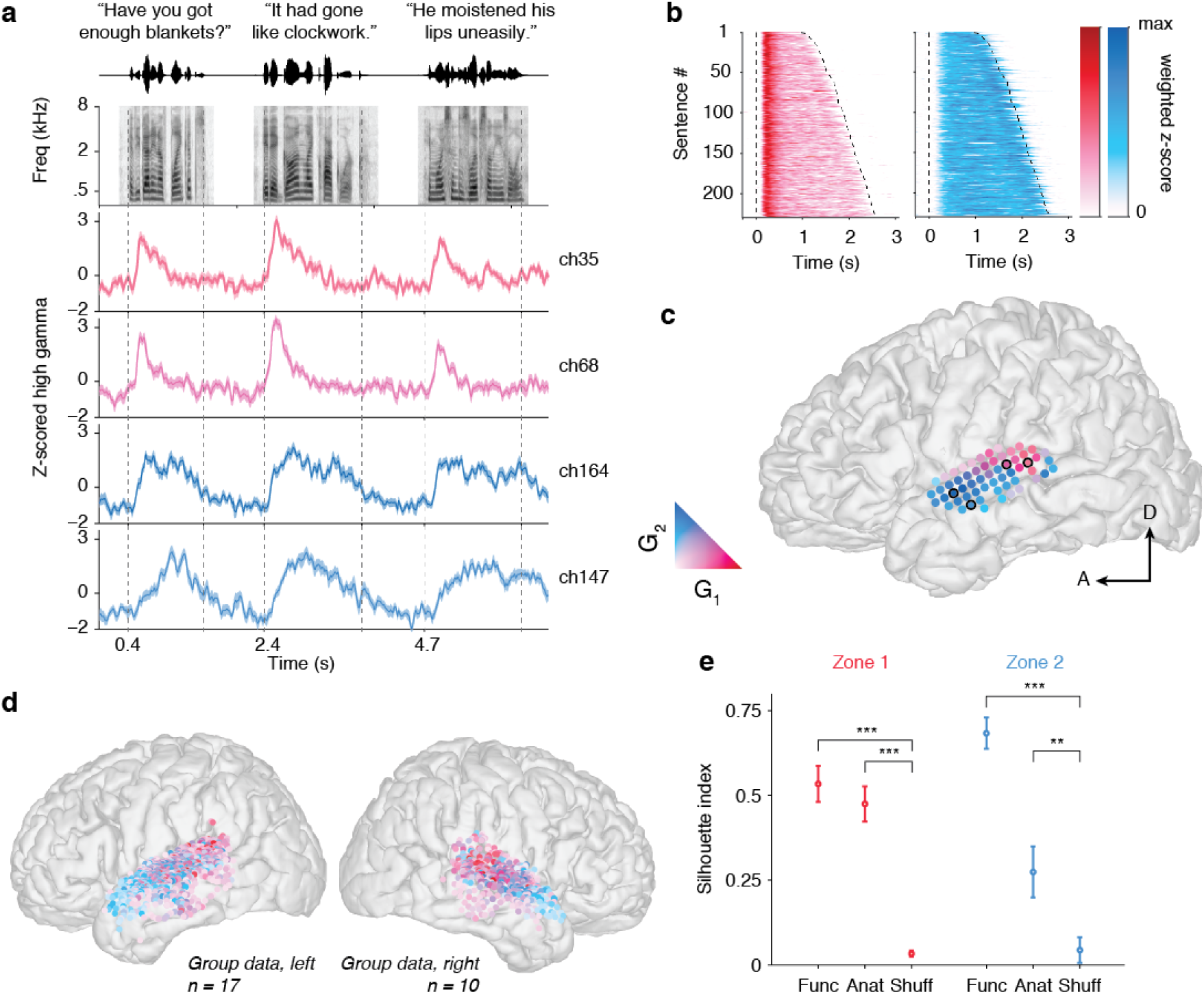
The superior temporal gyrus and middle temporal gyrus can be split into two spatially-segregated regions showing differential temporal responsivity to sentences. (a) Example responses to sentences from single electrodes in the first (red) and second (blue) clusters, colored according to their NMF activation weights. The waveform and spectrogram for each sentence is shown at the top. Zone 1 electrodes showed a strong response at sentence onset, sometimes followed by lower amplitude responses to other features within the sentence. Zone 2 electrodes showed differing responses throughout the sentence, and were not strongly selective for onsets. Sentence onsets and offsets are marked as dashed lines. (b) Average cluster time series across the Zone 1 and Zone 2 populations reveals that the two main distinguishing features of the neural response are (1) fast responses with strong activity at sentence onset and (2) slow responses, with weak responses at sentence onset and more sustained activity throughout the sentence. Each subplot shows the responses to all overlapping sentences across all subjects projected onto the NMF bases and sorted by sentence length. Sentence onset and offset are marked by dashed lines. (c) NMF activation weights on electrodes from one example left hemisphere subject, colored as in (a). Zone 1-type responses were observed in posterior STG close to the auditory core, whereas Zone 2 responses were found more anteriorly. Outlined electrodes identify the electrodes plotted in panel (a). (d) Activation weights for all left and right hemisphere subjects plotted on an average MNI brain. (e) Evaluation of functional and anatomical clustering goodness of fit using the silhouette index. A silhouette index close to 1 indicates good clustering, where within cluster distances are small and across cluster distances are large. Functionally, both Zone 1 and Zone 2 electrodes show tight clustering that was significantly higher than chance (Zone 1: *p* = 5.4 × 10^−5^, Zone 2: *p* = 8.3 × 10^−6^, Wilcoxon signed rank test). Anatomically, Zone 1 electrodes are close to one another in space and tend to be far away from Zone 2 electrodes, as evidenced by a high silhouette index that was significantly greater than a null shuffled distribution (p = 2.8 × 10^−4^, Wilcoxon signed rank test). Zone 2 electrodes are still significantly anatomically clustered (p = 7.4 × 10^−3^, Wilcoxon signed rank test), though less so than the Zone 1 electrodes.

### 2.2 Acoustic Representations in Zone 1 and 2

We fit spectrotemporal receptive field (STRF) models to each electrode separately to determine which combinations of spectrotemporal acoustic features would strongly elicit neural responses from these areas. Both Zone 1 and Zone 2 electrodes were well described by these models (NMF weighted average: rZone1 = 0.35 and rZone2 = 0.30). The weighted average of STRFs projected onto Zone 1 and 2 is shown in Fig. 2a. Both zones exhibited variable spectral selectivity (narrow and broad tuning), and appeared to be integrating sound information over relatively long time scales (up to 600ms for excitatory and inhibitory response). However, their temporal response profiles were substantially different. Zone 1 electrodes had strong preference for silence followed by sound onset, consistent with the onset sensitivity we observed at the sentence level. The excitatory part of the STRF was short in duration, while preceded by a relatively long inhibitory period. In contrast, Zone 2 electrodes appeared to have a long duration excitatory component only. Fig. 2b shows STRFs from single electrodes in Zone 1 (top) and Zone 2 (bottom). Both show selectivity for low, mid, and high frequency content. Some Zone 1 electrodes exhibited broadband, non-selective onset responses, whereas Zone 2 electrodes showed more complex spectral tuning such as adjacent excitatory and inhibitory sidebands.

**Figure 2:**
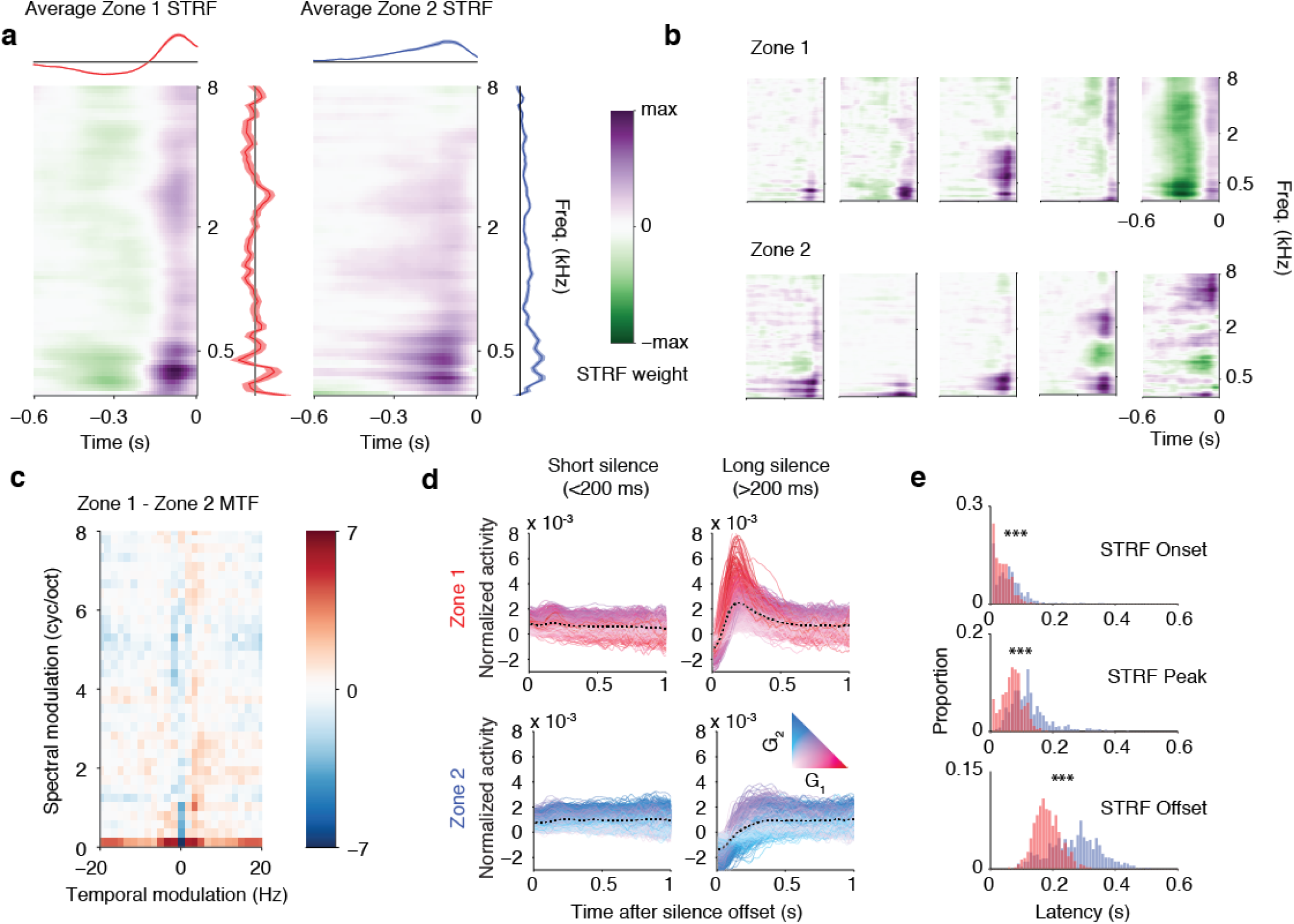
Zone 1 electrodes are onset detectors with selectivity to fast temporal modulations present in speech. Zone 2 electrodes are longer temporal integrators and are insensitive to speech onsets. (a) Zone 1 (left) and Zone 2 (right) weighted average spectrotemporal receptive fields (STRFs). Zone 1 electrodes show short latency responses and integrate over short temporal windows, whereas Zone 2 electrode responses are slower and longer. Temporal responses collapsed across frequency features are shown on the top, spectral responses collapsed across time are shown at right. (b) Example single electrode STRFs in Zone 1 (top) and Zone 2 (bottom). Overall, Zone 1 and 2 electrodes showed spectral selectivity over similar ranges, but differed in their temporal response profile. (c) Difference in average modulation transfer function across all Zone 1 and Zone 2 electrodes shows that Zone 1 electrodes show higher temporal modulation selectivity, whereas Zone 2 electrodes show higher spectral modulation selectivity for a limited range of modulation values. (d) Aligned responses to speech sounds after short (< 200 ms, left) and long (> 200 ms, right) silences. Zone 1 electrodes respond robustly after long silences, which can occur within a sentence or before sentence onset. Zone 2 electrodes respond to speech after short and long silences in a sustained manner. (e) Onset, peak, and offset response latencies calculated from Zone 1 and Zone 2 STRFs. Latencies were significantly greater in Zone 2 compared to Zone 1 electrodes (*p* < 0.001, Wilcoxon sign rank test), indicating longer temporal integration times for Zone 2.

While low level auditory areas perform a time-frequency decomposition of incoming sounds, higher auditory neurons are often sensitive to more complex combinations of features, including joint temporal and spectral amplitude changes or modulations (Eggermont, 2001; Shamma, 2001). Speech comprehension relies on encoding a relatively narrow set of spectral and temporal modulations within the spectra of all natural sounds (Elliott and Theunissen, 2009; Singh and Theunissen, 2003). To investigate this encoding in Zone 1 and Zone 2 electrodes, we measured selectivity to joint spectrotemporal modulations using the modulation transfer function (MTF), which describes whether these electrodes follow changes in spectral content, temporal content, or both (Singh and Theunissen, 2003). In agreement with previous work (Hullett et al., 2016; Pasley et al., 2012; Schönwiesner and Zatorre, 2009), we found higher temporal and lower spectral modulation selectivity in caudal Zone 1 electrodes, whereas rostral Zone 2 electrodes showed low temporal/high spectral modulation selectivity (MTF difference shown in Fig. 2c). The best temporal modulation was significantly greater in Zone 1 (mean ± stdev 1.5 ± 0.7 Hz) compared to Zone 2 (0.89 ± 0.9 Hz, comparison *p* < 0.001 Wilcoxon rank sum test). The best spectral modulation was significantly greater in Zone 2 (0.06 ± 0.2 cyc./oct.) compared to Zone 1 (0.03 ± 0.1 cyc./oct., p < 0.001, Wilcoxon rank sum test). The spectral modulation difference was strongest for low spectral modulations (< 1 cyc./oct.) that are critical for speech comprehension (Elliott and Theunissen, 2009).

In addition to sentence onsets, we were interested to know if similar responses were found within sentences after naturally occurring silent pauses between phrases. We plotted time-aligned Zone 1 and 2 responses to speech following short (< 200 ms) or long (> 200 ms) silences (Fig. 2d). Strong onset responses seen in Zone 1 occurred only after longer silences, consistent with the STRFs above, whereas Zone 2 exhibited slower, more sustained responses. Longer silences usually appeared at phrase boundaries in natural speech, and these findings suggest a similar response profile that strongly encodes these temporal landmarks in speech. These onset responses were observed even when stimuli were played backwards or were spectrally rotated to remove phonetic and lexical content (Supplemental Fig. 4). Thus, although by analyzing modulation selectivity we were able to replicate previous findings of high temporal modulation representation in pSTG, the most parsimonious explanation of these data appears to be that Zone 1 regions are onset selective, rather than simply selective for high temporal modulations. This also explains our previous results whereby modulated ripple noises with spectral and temporal modulation content did not elicit strong activity in STG beyond an initial onset response (Hullett et al., 2016).

The temporal integration profiles of brain areas can provide insights as to the type of information that is being encoded. To quantify the difference in latencies and temporal integration profile in Zone 1 and 2, we used the STRF for each electrode to derive response latencies; we calculated the onset, peak, and offset of the excitatory component in each STRF. Zone 1 electrodes exhibited earlier onset, peak, and offset latencies compared to electrodes in Zone 2 (p < 0.001, Wilcoxon rank-sum test; Fig. 2e), but that the difference was most pronounced at offset, where the difference in average offset latencies was 76 ms. The mean duration of the excitatory response was 139 ± 1 ms in Zone 1, and 196 ± 2 ms in Zone 2. These temporal properties are consistent with previous findings (Honey et al., 2012; Lerner et al., 2011; Nourski et al., 2014) and have been interpreted as evidence of serial processing within the “ventral stream”, with the idea that posterior areas are lower order. However, the results here suggest that the representations are fundamentally different between the zones, suggesting more of a parallel operation. First, many Zone 1 electrodes are non-selective in the spectral domain, whereas Zone 2 responds to spectrally complex sounds. Second, if Zone 1 were low-level, it would be activated throughout the sentence, not primarily driven by the onset. Third, and most importantly, how the two zones integrate sound information over time is completely different.

In sum, our acoustic analysis indicated that Zone 1 electrodes are onset detectors with varied spectral selectivity and fast temporal modulation selectivity, while Zone 2 electrodes are long temporal integrators that are not sensitive to onsets and encode spectral modulations important for speech comprehension. Next, we wanted to examine how these acoustic properties related to phoneme feature representations in each zone.

### 2.3 Local phoneme feature embedding in Zone 1 and Zone 2

Phonetic features refer to articulatory cues in speech, such as plosive, nasal, fricative, and vowel, which describe how the sounds that define different categories of phonemes are produced by the vocal tract (Ladefoged and Johnson, 2011). Previously we showed that the human STG exhibited selectivity for phonetic features (Mesgarani et al., 2014), however no consistent spatial map for these features was found across subjects.

We speculated that plosives characterized by silence followed by broadband burst (e.g. /ba/, /pa/, and /ta/) might be selectively processed in Zone 1 because of its sensitivity to temporal modulation information. Conversely, we predicted that Zone 2 would be sensitive to spectral modulation content in vowels. As before, we fit a linear model to predict electrode activity, this time employing a reduced binary feature matrix to represent the presence or absence of phonetic features (vowels, nasals, fricatives, and plosives) in the sentence stimuli (see Methods for details). In addition, we added a feature identifying the start of each sentence to better model onset responses, and we added a speech vs. silence term to model overall increases in activity during speech sounds.

**Figure 3:**
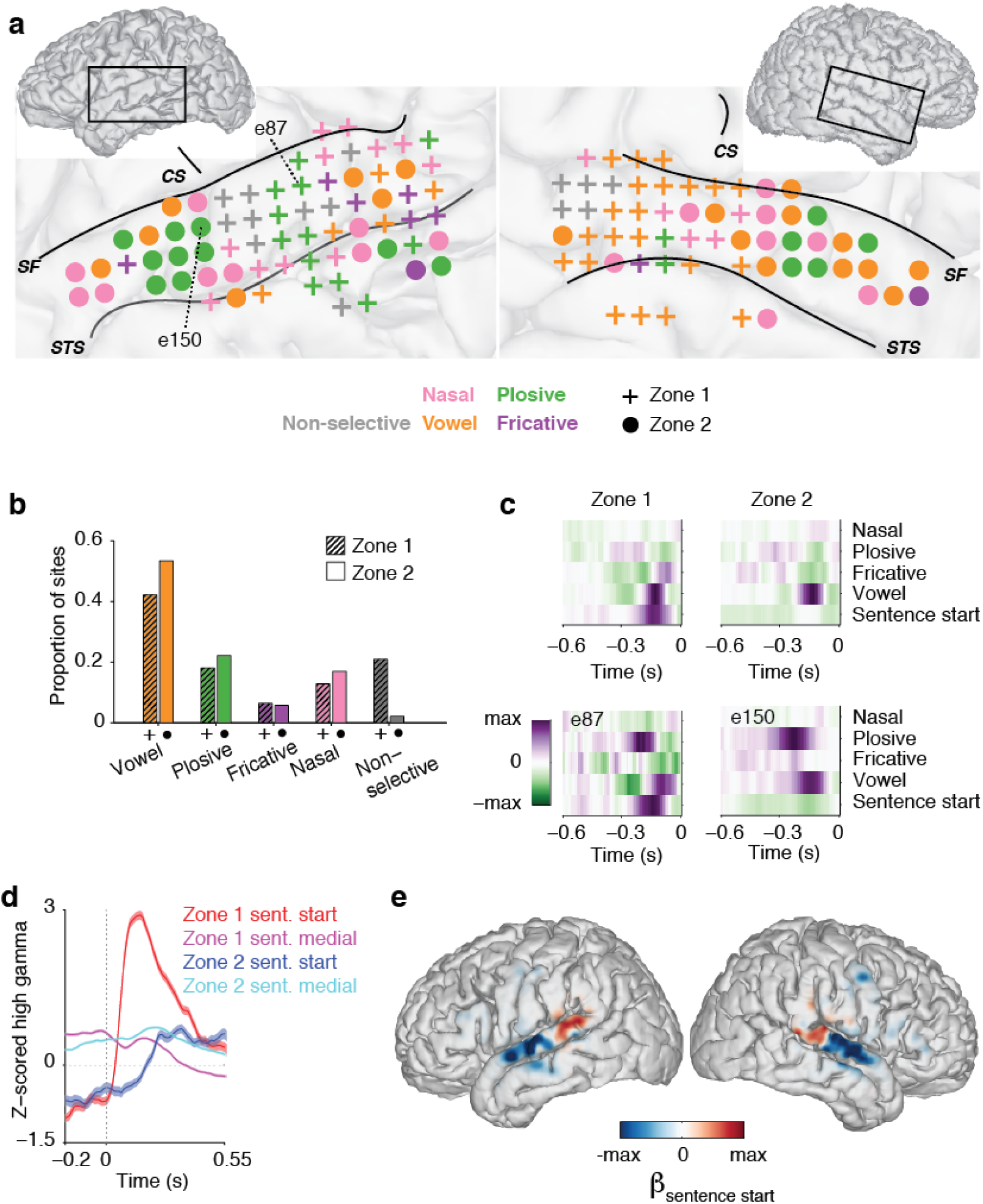
Zone 1 and Zone 2 electrodes exhibit overlapping phonetic feature selectivity. (a) Example phonetic feature maps for two subjects (EC36, left and EC28, right). Electrodes are colored according to their phonetic feature selectivity (non-selective, nasal, vowel, plosive, or fricative). Zone 1 electrodes are denoted by the + symbol, Zone 2 electrodes by a circle. SF=Sylvian fissure, STS = superior temporal sulcus, CS = central sulcus. (b) The distribution of phonetic class selectivity is similar across Zone 1 and Zone 2 fields (*p* = 0.2, Chi square test). The proportion of sites exhibiting selectivity for each of the five phonetic features is shown for Zone 1 (dark bars, left) and Zone 2 (light bars, right) sites. (c) Example electrodes with similar phonetic feature selectivity (top: vowels, bottom: plosives) belonging to the Zone 1 (left) and Zone 2 (right) classes. Note that plosive-selective electrodes often show selectivity to plosives followed by a vowel. (d) Example aligned high gamma responses to plosive phonemes at sentence start and sentence medially for electrodes shown in bottom left and right of panel (c). The Zone 1 electrode shows strong modulation by phoneme position, whereas the Zone 2 electrode does not. Because both sets of responses were Z-scored relative to the average activity across the entire experiment, the sentence-initial plosives begin with a baseline below zero, whereas activity associated with mid-sentence plosives may also contain responses to other phonemes preceding the sound of interest. Traces show mean ± standard error across trials. (e) Sentence start weights for all subjects, plotted as a heat map on the CVS atlas brain. Positive weights (indicating increased response to sentence start) are shown in red and are strongly localized to the posterior STG in both left and right hemispheres.

Contrary to our expectations, we observed overlapping phonetic feature representation in Zone 1 and Zone 2 – that is, we did not find segregation of consonants and vowels in these areas. Instead, we found evidence of single electrode selectivity for all phonetic feature classes: vowels, nasals, fricatives, and plosives, in both Zone 1 and Zone 2. Fig. 3a illustrates this diversity of phonetic feature representation in two example subjects, with Zone 1 electrodes demarcated by + and Zone 2 electrodes as circles. For example, both Zone 1 electrode 87 and Zone 2 electrode 150 in panel (a) show responses to plosives (green). “Nasal”, “fricative”, “plosive”, or “vowel” selective electrodes can be observed in both Zone 1 and 2 in the left and right hemispheres. The proportion of sites tuned to a particular phonetic feature did not significantly differ across Zone 1 and 2 (p=0.2, Chi-square test; see Fig. 3b for the proportion of sites across all subjects that exhibited selectivity for each feature). This unexpected result is perhaps reconciled by the observation that the STRF temporal integration time was substantially longer than the length of a phoneme (in TIMIT, the average labeled phoneme length is 80.2 ± 5.8 ms (mean ± stdev), whereas our STRF integration times were much longer (in Zone 2, many responses persisted through almost the entire 600-ms STRF delay period).

Examples of the feature-temporal receptive fields for four electrodes are shown in Fig. 3c. The top two panels show a Zone 1 “vowel” electrode and a Zone 2 “vowel” electrode, whereas the bottom panels show a Zone 1 “plosive” and a Zone 2 “plosive” electrode (also shown on the brain in Fig. 3a). Interestingly, many “plosive” selective electrodes in both Zone 1 and Zone 2 were sensitive to the sequential combination of plosive followed by vowel. Within Zone 1, some electrodes responded strongly to specific phonetic features, with a higher response at the start of a sentence (as evidenced by high weights on the “sentence start” feature, see Fig. 3d for raw example responses to plosives at sentence start and sentence-medially). Therefore, response selectivity to phonetic features is jointly encoded with the sentence start feature.

This strong differentiation of sentence-initial vs. sentence-medial phoneme feature representation for Zone 1 electrodes was strongly spatially localized. Electrodes with a high response at sentence start were located exclusively in the posterior STG (Fig. 3e), in a distinct region largely overlapping with Zone 1. Onset weights were strongly positively correlated with Zone 1 NMF weight (Spearman rho = 0.62) and strongly negatively correlated with Zone 2 NMF weights (Spearman rho = −0.75). In contrast, we found no evidence for a reliable or consistent spatial map of phonetic feature selectivity (Supplemental Fig. 5).

Altogether, these results demonstrate that phonetic feature encoding is differentiated at the local-scale of individual electrodes, whereas the temporal parameters (“onset” vs. “sustained”) are a more global-scale organizational property that partitions the STG.

### 2.4 Tonic syllable encoding in the Zone 2 cortex

We next wanted to better understand what kind of acoustic or linguistic properties were being processed in Zone 2. Given the longer temporal integration there, we evaluated whether Zone 2 electrodes were sensitive to suprasegmental features, such as stress patterns. In particular, we were interested in an intonational cue, called the “tonic” syllable, which usually falls on the last stressed syllable of a sentence. There is one tonic syllable for every tone unit, which begins the terminal prosodic contour, such as the pitch rise of certain questions, or the pitch drop that conveys finality in declarative sentences (Halliday, 1963; Ladefoged and Johnson, 2011). For example, in the sentence “It was nobody’s fault” (Fig. 4a), both the vowels /ow/ in “nobody” and the /a/ in “fault” are stressed, but /a/ is more important intonationally.

In our encoding model for phonetic features, we marked vowel stress as well as the tonic syllable in each sentence. Both Zone 1 and Zone 2 encoded vowel stress, with larger responses for high stress vowels (Fig. 4b). However, a subset of Zone 2 electrodes showed a prominent response to the tonic syllable, above and beyond the responses for stressed syllables (Fig. 4b). By contrast, Zone 1 electrodes responded to the tonic syllable only as another stressed syllable, apparently insensitive to the intonational aspect. This suggests that Zone 2, with its longer temporal integration period and selectivity for spectral modulations, also preferentially responds to prosodic features in the sentence that are important for linguistic meaning.

A single electrode example of this response type is shown in Fig. 4c. The spectrogram for the sentence “It was nobody’s fault” is shown again at the top, with the amplitude contour (sum of the spectral power across frequencies) in pink and the F0 pitch contour in blue. Below, the response to this sentence is shown for the electrode depicted in Fig. 4d (also see Supplemental Fig. 6). The degree of stress for each vowel portion of the syllable is shown in pink, with darker pink depicting high stress. The individual regression weights for this electrode are shown in Fig. 4e, with a strong response to high stress syllables, and an additional increase in activity if the syllable was also the tonic syllable. When data was high-pass filtered to remove the contribution of the sentence envelope (see Methods), NMF basis functions showed this strong modulation by the tonic syllable for Zone 2 compared to Zone 1 (Fig. 4f). Zone 2 had a strong increase in cortical activity locked to the tonic syllable (at t = 0) before the sentence offset, whereas Zone 1 was still most responsive at sentence onset.

Since the tonic syllable is defined using linguistic criteria, we evaluated whether tonic syllable responses were simply an effect of acoustic differences compared to other syllables. By definition, the tonic syllable does not necessarily need to be the loudest syllable or the syllable with the highest degree of pitch-change (a strong indicator of intonational stress). We computed the average spectral power as the sum of power in the spectrogram across all frequencies during unstressed, low stress, and high stress syllables, as well as the tonic syllable. In addition, we computed the average change in pitch during these syllables. The tonic syllable overlapped substantially in these measures compared to other syllable types (Fig. 4g), so it is unlikely that these effects are driven by overall acoustic differences in the tonic compared to other syllables. Instead, we posit that the tonic syllable is a critical temporal landmark in continuous speech, as posited by linguists to mark a key moment of informational delivery in the prosodic contour (Hultzén, 1959).

**Figure 4:**
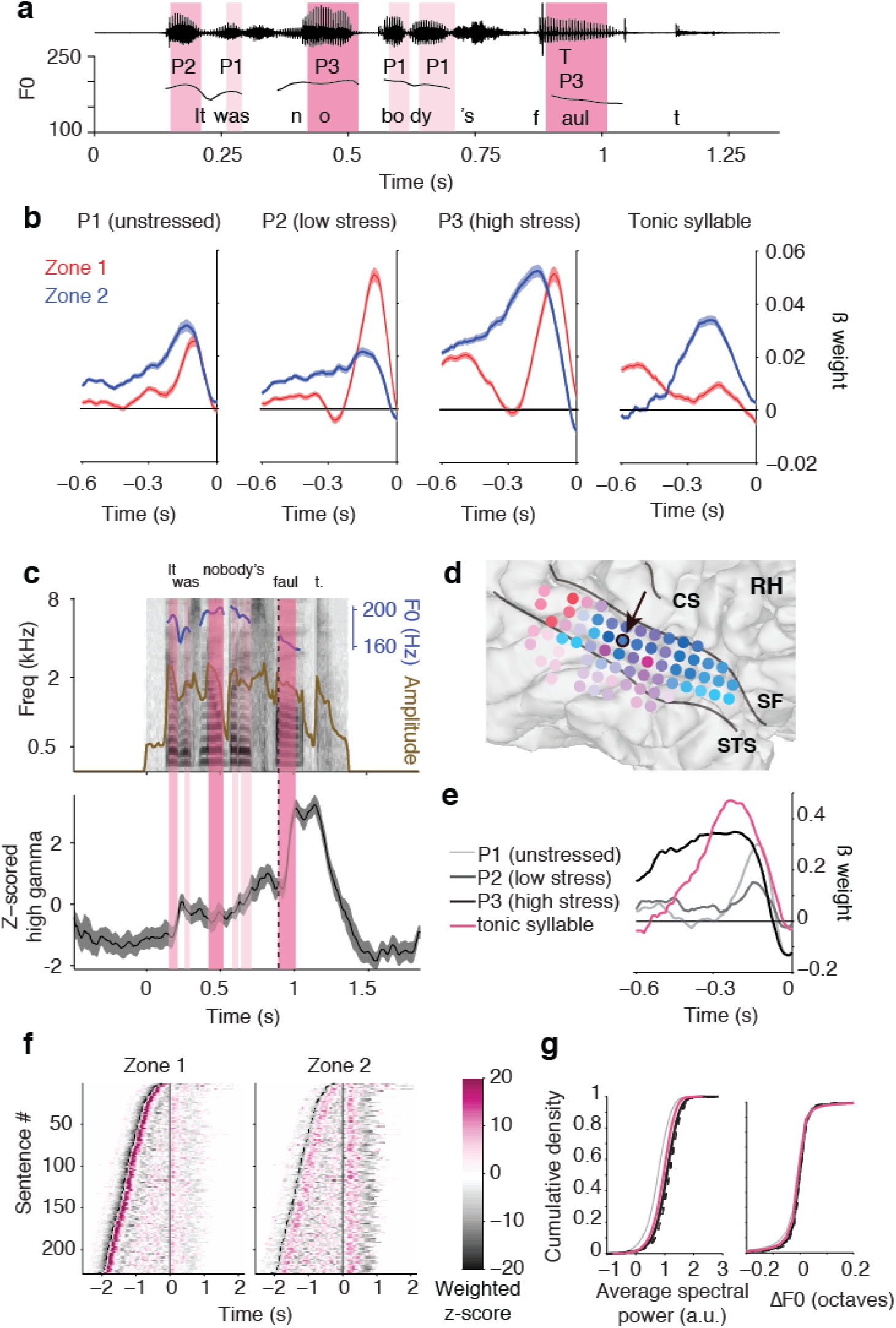
Zone 2 electrodes show enhanced response to the tonic syllable, the most significant syllable in the tone unit that carries important linguistic content. (a) Example sentence waveform (top), annotated for stress (P1-P3), tonic syllable (T), and pitch contour (bottom). Here the tonic syllable, the /a/ in “fault”, does not show a higher amplitude nor stronger pitch contour, but rather gives the sentence its meaning and serves as the most linguistically important syllable in the sentence. (b) Average regression weights across Zone 1 (red) and Zone 2 (blue) for unstressed (P1), low stress (P2), and high stress (P3) vowel components of the syllable, as well as the response to the tonic syllable. In general, responses in both sets of electrodes increase with increasing stress. Responses in posterior Zone 1 lead those in anterior Zone 2. In Zone 2 electrodes only, there is an enhanced response for the tonic syllable above and beyond what is predicted by stress alone. (c) Example response from a single electrode in Zone 2 to the tonic syllable (dashed line) in the sentence “It was nobody’s fault”, with stress indicated as in (a) by pink shading. (d) Location of electrode in panel (c) is shown outlined in black on zoomed-in right hemisphere patient’s brain. Other electrodes are colored according to their weight on Zone 1 (red) and Zone 2 (blue). CS = central sulcus, SF = Sylvian fissure, STS = superior temporal sulcus. (e) Regression weights for this electrode. The electrode responds strongly to high stress vowels (P3), but also shows an enhanced response to the tonic syllable (pink), which can co-occur with P3 and P2 syllable types. (f) Tonic syllable-aligned responses projected onto Zone 1 and Zone 2, high-pass filtered at 1 Hz to remove overall sentence envelope fluctuations. First dashed line indicates sentence onset; solid line indicates the onset of the tonic syllable (t=0). Zone 2 electrodes show strong time-locked responses to this syllable. (g) Enhanced responses to tonic syllable cannot be explained by increased loudness or pitch changes during this syllable. Left panel shows cumulative density function over the average spectral power (the sum across all frequency bands in the spectrogram) within each syllable type. The tonic syllable showed significantly lower average spectral power compared to stressed syllables (P2 and P3) and significantly higher average spectral power compared to unstressed (P1). Right panel shows cumulative density plot of change in fundamental frequency (pitch change) across syllable types. The tonic syllable showed a medium degree of pitch change compared to other syllable types. P2 and P3 distributions are shown including (solid lines) and excluding (dashed lines) tonic syllable, since the tonic syllable was a subset of these categories.

### 2.5 Zone 1 and 2 response types partition the entire auditory-responsive speech cortex

The exposed lateral STG is only a portion of the human auditory cortex, and is an extension of auditory processing in “core” and “belt” areas of the temporal plane (Hackett et al., 2014; Kaas and Hackett, 2000; Moerel et al., 2014). Other groups have successfully employed depth recording electrodes in Heschl’s gyrus (HG) to show that core auditory cortical areas show tonotopic organization, fast latency responses to click trains, and can track pitch changes in pure tones (Brugge et al., 2009; Griffiths et al., 2010; Howard III et al., 1996; Steinschneider et al., 2014). In four participants, we recorded simultaneously from the superior temporal plane and the lateral surface with high-density grids (4 mm center-to-center spacing, see example of placement on the temporal lobe for one subject in Fig. 5a)–thereby covering the entire auditory cortical pathway. For these participants, microdissection of the Sylvian fissure was clinically indicated for accessing circumscribed insular tumors during awake craniotomy. This allowed us to temporarily access the core primary auditory cortex, including HG and nearby belt areas in the planum temporale (PT) and planum polare (PP). Electrodes on the HG and areas of PT responded robustly to speech (Fig. 5b) and to pure tones (Fig. 5c). As expected, in the posteromedial portion of HG we found clear evidence for tonotopic progression from low to high frequency tuning, using pure tone and speech stimuli. In contrast, speech responsive electrodes in the STG (bottom row electrodes in Fig. 5b and 5c) were often spectrally multi-peaked, broadly tuned or not responsive to pure tones, and therefore were not amenable to address tonotopy.

With NMF clustering including these electrodes, we found that Zone 1 and 2 response profiles observed in the STG were recapitulated in the temporal plane. They followed the same caudal/rostral distinction as the responses across the lateral STG surface (Fig. 5d). Electrodes over the posteromedial HG and PT mostly belonged to Zone 1, whereas rostral PP electrodes projected strongly onto Zone 2 (see Fig. 5e for example single electrode responses to sentences as well as their STRFs). HG and PT electrodes showed strong onset responses and short temporal integration times, whereas PP electrodes showed slower responses similar to rostral STG. In contrast to the Zone 1 electrodes in STG, however, Zone 1 electrodes in HG did not show responses confined to sentence onsets, and instead showed fast, phasic onset responses followed by additional responses to features throughout the sentence. Electrodes in PP and rostral temporal plane exhibited more spectral broadband tuning, unlike the stereotypic narrow, highly tuned responses in the core. The average STRF for Zone 1 and 2 electrodes on the temporal plane (Fig. 5f) were very similar in appearance to the average STRFs for Zone 1 and 2 on the lateral surface (Fig. 2a), again exhibiting fast, onset-selective responses in Zone 1, and longer temporal integration in Zone 2. The temporal integration window, calculated as the difference in STRF offset and onset latencies, was significantly shorter for temporal plane electrodes in Zone 2 (0.20 ± 0.05 s for temporal plane, 0.23 ± 0.07 s for STG/MTG, *p* = 0.03 Wilcoxon rank sum test) and at trend level in Zone 1 (0.15 ± 0.04 s for temporal plane, 0.16 ± 0.04 STG/MTG, *p* = 0.14, Wilcoxon rank sum test), indicating a medial to lateral increase in temporal integration in addition to the aforementioned caudal to rostral differences.

Although temporal plane electrodes responded robustly to speech features, they were generally better fit by a spectrotemporal than a phoneme feature model in both Zone 1 and Zone 2 (average *r*_spectrotemporal_ =0.31±0.18, *r*_feature_ = 0.29±0.17, *p* = 7.8 × 10^-9^, Wilcoxon signed rank test). In contrast, the phoneme feature model outperformed the spectrotemporal model in STG and MTG sites *r*_spectrotemporal_ = 0.28 ± 0.15, *r*_feature_ = 0.30 ± 0.16, *p* = 1.9 × 10^-66^, Wilcoxon signed rank test), suggesting that some acoustic to phonetic transformation occurs in lateral parabelt regions rather than the temporal plane.

Finally, work by our group (Cheung et al., 2016) and others (Cogan et al., 2014; Wilson et al., 2004) has shown responses to speech during passive listening in the sensorimotor cortex as well as inferior frontal gyrus. When including these electrodes, we again found a separation of responses into Zone 1 and 2 response types (Supplemental Fig. 7a). Although the overall magnitude of the cluster weights in this area was lower than in the classical auditory areas, the response profile for single electrodes within these regions was similar, as was the overall structure of the STRFs (Supplemental Fig. 7b-c). These results suggest that the Zone 1 and Zone 2 distinctions not only apply to the parabelt STG, but also appear to be a fundamental organizing response feature across the entire auditory-responsive speech cortex.

**Figure 5:**
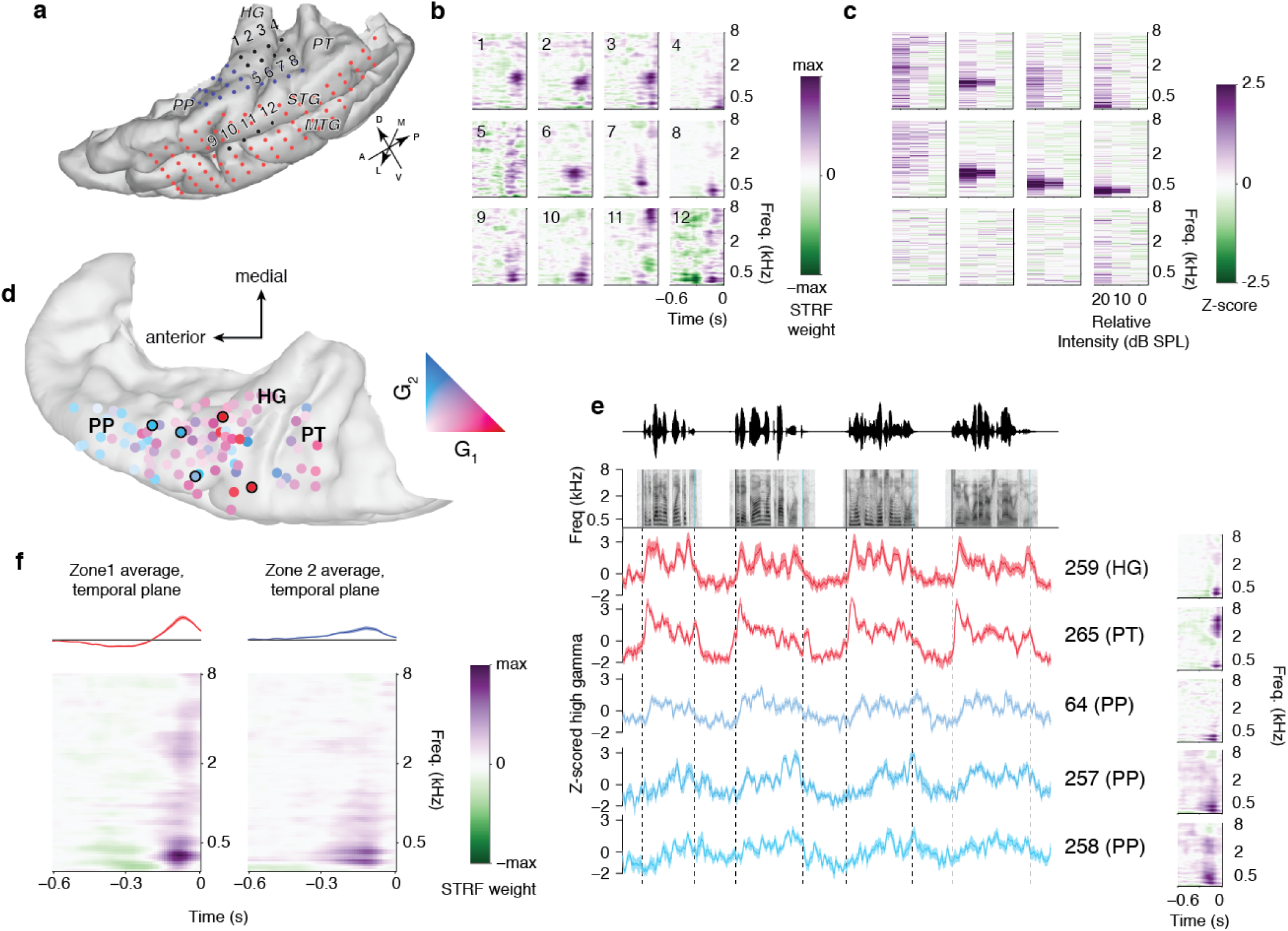
Zone 1 and 2 are an organizing feature across the entire auditory hierarchy. (a) Temporal lobe reconstruction for one patient with one grid on the temporal plane (blue) and another on the lateral surface including STG (red). Numbered electrodes on the Heschl’s gyrus (HG), planum temporale (PT), and superior temporal gyrus (STG) match receptive fields in (b) and (c). (b) Example spectrotemporal receptive fields (STRFs) for temporal plane and STG electrodes. Electrodes on HG, PT, and STG showed robust responses to speech. On the temporal plane grid, we saw evidence of tonotopic organization in some electrodes; this same organization was not observed in the STG. (c) Responses to pure tones for the same electrodes shown in (b). HG and PT showed robust, narrowly tuned tone responses. The STG showed either broadband, high threshold tone responses (bottom right), or no discernible tone response. (d) All temporal plane electrodes warped and plotted on an MNI brain (cvs_avg35_inMNI152 atlas). Electrodes are colored according to their weight on Zone 1 (red) or Zone 2 (blue), as in Figure 1. Zone 1 electrodes were localized to the posterior temporal plane, including PT and HG, whereas responses in PP mostly projected onto Zone 2. (e) Single sentence responses to four sentences from electrodes outlined in (d), and their corresponding STRF at right. Zone 1 electrodes in HG and PT showed narrow temporal selectivity and strong responses at sentence onset, whereas Zone 2 electrodes in PP showed broader, later responses. (f) Average STRF for all Zone 1 (left) and Zone 2 (right) electrodes on the temporal plane. Zone 1 electrodes in this region also showed a characteristic onset-like response, whereas Zone 2 electrodes were long temporal integrators.

### 2.6 Parallel streams encode the temporal landmarks in continuous speech dynamics

We identified distinct pathways of the speech-responsive cortex that appear to respond differentially to onset and non-onset components of speech and that integrate over short and long time scales. These observations suggest that the auditory system is highly sensitive to the temporal dynamics intrinsic to the overall structure of phrases and sentences. Thus, to visualize how Zone 1 and 2 populations contribute to the temporal dynamics of natural speech processing, we performed a cortical state-space analysis (Briggman et al., 2005; Churchland et al., 2012; Kao et al., 2015; Mazor and Laurent, 2005) in which responses to sentences were projected onto NMF components. We first examined the relationship between Zone 1 and Zone 2 responses across all anatomical sites (Fig. 6a, also see Supplemental Fig. 8). Immediately following sentence onset, the state space trajectory was dominated by Zone 1 responses, then moved towards Zone 2 response types. The neural trajectories eventually converged on a relatively confined region of the state-space representing the tonic syllable (pink) before the sentence end. For single sentences, the trajectories through this state space show a stereotyped response profile. In Fig. 6b at left, the sentence “But why pay her bills?” shows a sweep into the onset Zone 1, followed by coactivation of Zone 2 and an additional increase in activity in Zone 2 during the tonic syllable (the /er/ in “her”, pink), followed by a decline in activity toward the end of the sentence after the tonic syllable. At right, a natural sentence containing a substantial pause in the middle (“Then he – then what?”) results in two rotations through this state space (“then he” in yellow hues, “then what?” in yellow/orange hues). Notably, when marking the position of other phonetic features (for example, vowels, fricatives, or plosives) within this state space, the phonetic features did not occupy a defined region of the global dynamics represented here, but rather were distributed throughout the sentence trajectory (Fig.6c–e).

**Figure 6:**
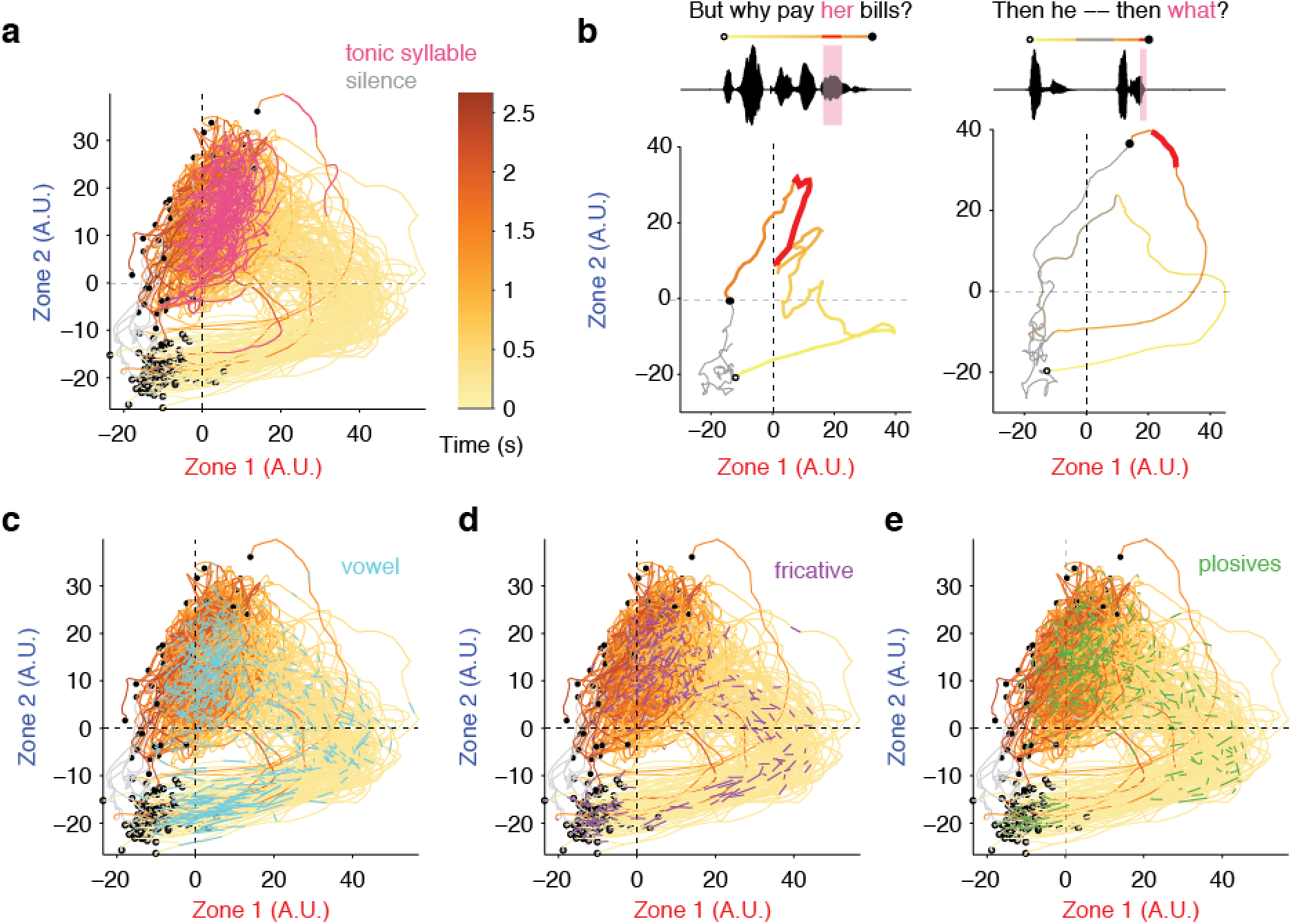
State space trajectories for Zone 1 and Zone 2 electrodes encode temporal landmarks in speech. (a)State-space representation for all electrodes (*n* = 2,100) projected onto Zone 1 and Zone 2 during single sentences. Each trajectory within the space represents one sentence. Each sentence starts and ends with silence (marked in gray). The open circle indicates speech onset, and is followed by a colored line that progresses from light yellow to orange as time progresses in the sentence. The tonic syllable is marked on the trajectory in pink. Sentence offset is marked as the filled circle. Across all sentences, neural activity follows a stereotyped sweep through Zone 1, then coactivating Zone 2. The tonic syllable occupies a distinct area of this cortical state space, as seen in previous analyses, where it projects highly onto Zone 2. (b) Single sentence trajectories for a short sentence (left) and a sentence broken by a brief pause (right). In the case of a pause, neural activity traverses the space twice – once for each segment of the sentence. (c-e) Responses to phonetic features are widely distributed within global state space trajectories. State space trajectories are marked for the presence of vowels (c), fricatives (d), and plosives (e), which, unlike the tonic syllable, occupy multiple regions of the cortical state space.

## 3 Discussion

Unsupervised clustering of human cortical responses to speech identified two distinct regions that were remarkably consistent across 27 participants. These zones appear to represent both low-level features found throughout the auditory system, such as onsets, and high-level features which have not been previously observed, such as the tonic syllable. Together, our findings demonstrate a potential neural code for demarcating the timing or position of linguistically important events in natural speech, thereby providing contextual information to the segmental phonetic feature processing locally within each large region.

Our work uncovered two major response types in Zone 1 and Zone 2 with “transient” and “sustained” responsivity to sentences. Such “phasic” and “tonic” response types have been observed in single unit electrophysiological recordings throughout the auditory system, including the temporal lobe auditory cortex (Eggermont, 2001; Malone et al., 2015; Romanski et al., 1999), auditory brain stem (Atencio et al., 2012; Zheng and Escab, 2008), and even prefrontal cortex, where they may be involved in decision making and object identification within the “ventral stream”(Bizley and Cohen, 2013; Romanski and Goldman-Rakic, 2002; Romanski et al., 1999; Tsunada et al., 2016). Previous studies, however, have not clearly documented the spatial segregation of these response types in auditory cortex. As a result, it was a surprise to discover that such response distinctions can be consistently regionalized at a macro-anatomical level. To our knowledge only limited fMRI evidence has suggested transient and sustained cortical responses in humans (Harms and Melcher, 2003; Seifritz et al., 2002; Werner and Noppeney, 2011), although those responses were over much longer timescales (seconds and minutes), largely confined to the temporal plane, and in context of synthetic stimuli or scanner noise rather than natural speech.

Many previous studies, including our own, may have overlooked these properties in search of more canonical acoustic and speech features, such as phonemes and syllables, in the human STG. Here, the data-driven approach combined with large-scale coverage, dense sampling, and real-time ECoG recordings contributed directly to the novel functional clustering observed here. Certainly, a model-based approach would have found clear neural correlates to sentence onsets and tonic syllables had they been sought out.

Despite the limited previous demonstration of functional organization, substantial anatomical evidence exists for differentiating the caudal and rostral auditory cortex (Kaas and Hackett, 1999, 2000; Rauschecker and Scott, 2009; Rauschecker et al., 1997; Romanski et al., 1999). We refer to our caudal and rostral areas as Zone 1 and Zone 2, respectively, after anatomical studies showing parallel topographic projections from the inferior colliculus to the medial geniculate body (Cant and Benson, 2007). These streams appear to persist up to the primary auditory, belt, and parabelt areas, and may be part of the “where” (caudal) and “what” (rostral) pathways described in non-human primates (Bizley and Cohen, 2013; Kaas and Hackett, 1999; Rauschecker, 2012; Romanski et al., 1999). Our results from temporal plane and lateral STG support connectivity studies in non-human primates that posit two dimensions of auditory processing along the medial-lateral and caudal-rostral axes, respectively (Hackett, 2011). Furthermore, core and planum temporale (including the caudal parabelt) are distinguished by their heavy cortical myelination patterns (Dick et al., 2012; Glasser and Van Essen, 2011; Hackett et al., 2001), which is likely to support the rapid temporal processing in these areas. As for the “dorsal” vs. “ventral” stream model (Hickok and Poeppel, 2007), we believe this to be a separate distinction. In our data, the distinct Zone 1 vs. Zone 2 responders were found within the ventral stream regions of STG and MTG. Dorsal stream sites (e.g. motor cortex, supramarginal gyrus) can also load as Zone 1 or Zone 2 types, but typically less strongly than in ventral regions.

Our results suggest that this parcellation is a fundamental aspect of auditory cortex organization and is likely not specific to speech processing (Steinschneider et al., 2013). Nevertheless, these basic response properties have implications for detecting the timing of linguistically important events. A major novel finding of the present study is that the “tonic syllable”(Halliday, 1963) in linguistic prosody elicits a strong, selective response from a subset of Zone 2 electrodes, consistent with their proximity to known pitch processing areas (i.e. the primate “pitch center”(Bendor and Wang, 2005)). Sensitivity to onsets can play a critical role in parsing sentence and phrase boundaries using acoustic cues, in combination with other high-level syntactic cues (Ding et al., 2015). The onset detectors observed in Zone 1 may be critical for auditory scene analysis (Bregman, 1994), since spectral energy from single sources are generally temporally coherent (O’Sullivan et al., 2015; Shamma et al., 2011). We observed very few responses to offsets compared to onsets, in accordance with neurophysiological and behavioral evidence that sound onsets are given greater perceptual weight (Phillips et al., 2002).

We have described here a major division of the entire auditory cortical system that supports multiple levels of spectrotemporal, phonological, and linguistic representations. This defining property of auditory cortical organization suggests how neural populations combine dynamically to support speech perception, and likely reflects a general mechanism for processing natural sounds. While these findings demonstrate the extraction of multiple dimensions of acoustic-phonetic and temporal cues in speech, a major challenge is to understand how such information is fully integrated at higher cortical levels to support language comprehension.

## 4 Methods

### 4.1 Participants

Participants included 27 patients (13M/14F, age: 36 ± 12 years) implanted with high-density subdural intracranial electrode grids (AdTech or Integra, 256 channels, 4mm center-to-center spacing and 1.17mm diameter) either chronically as part of their clinical evaluation for epilepsy surgery, or in an acute intraoperative setting for tumor resection. All procedures were approved by the University of California, San Francisco Institutional Review Board, and all patients provided informed written consent to participate. 17 subjects were implanted with left hemisphere grids, and 10 subjects were implanted with right hemisphere grids. Details of implantation (hemisphere, sex, handedness, language dominance, and seizure focus) are in Supplementary Table 1. In the 4 patients with tumors necessitating dissection of the Sylvian fissure and exposure of the temporal plane, we also acquired neural recordings directly from the temporal plane using 32 channel (8 x 4) or 64 channel (8 x 8) grids with similar specifications to the lateral grids (Integra, 4mm center-to-center spacing and 1.17mm diameter). For these patients, recordings were acquired simultaneously from the temporal plane and the lateral surface of the brain (including STG/MTG).

### 4.2 Neural recordings

Electrophysiological recordings were acquired at a sampling rate of 3051.8 Hz using a 256-channel PZ2 amplifier or 512-channel PZ5 amplifier connected to an RZ2 digital acquisition system (Tucker-Davis Technologies, Alachua, FL, USA). We recorded the local field potential from each electrode, notch-filtered the signal at 60 Hz and harmonics (120 Hz and 180 Hz) to reduce line-noise related artifacts, and re-referenced to the common average across channels sharing the same connector to the preamplifier (Cheung et al., 2016). We then used the log-analytic amplitude of the Hilbert transform to bandpass signals in the high gamma range (70-150 Hz), using 8 logarithmically-spaced center frequency bands and taking using first principal component across these bands to extract stimulus-related neural activity (Edwards et al., 2009; Moses et al., 2016; Ray and Maunsell, 2011). High gamma signals were then downsampled to 100 Hz for further analysis. Signals were z-scored relative to the mean and standard deviation of activity across each recording session.

### 4.3 Cortical surface extraction and electrode visualization

We localized electrodes on each individual’s brain by co-registering the preoperative T1 MRI with a postoperative CT scan containing the electrode locations, using a normalized mutual information routine in SPM12. For intraoperative patients for whom no CT scan was available (4 subjects), electrode localization was performed manually using intraoperative photographs of grid placement. Pial surface reconstructions were created using Freesurfer. For visualization of electrode coordinates in MNI space, we performed nonlinear surface registration using a spherical sulcal-based alignment in Freesurfer, aligning to the cvs_avg35_inMNI152 template (Fischl et al., 1999). This nonlinear alignment ensures that electrodes on a gyrus in the subject’s native space remain on the same gyrus in the atlas space, but does not maintain the geometry of the grid.

### 4.4 Stimuli

*TIMIT sentences*. Speech stimuli were 499 sentences taken from the TIMIT acoustic-phonetic corpus (Garofolo et al., 1993), spoken by 286 males and 116 females from different regions of the United States of America. Most sentences were repeated 1-2 times; for the majority of patients (23/27), a subset of 10 sentences was repeated 10-22 times. Stimuli were played through free-field speakers (Logitech), and presentation was controlled using custom MATLAB software on a Windows Laptop. Sentences were presented in pseudorandom order with 0.4 s of silence in between each.

*Pure tone stimuli*. For subjects with temporal plane coverage, we also played pure tone stimuli, synthesized as 50-ms duration 5-ms cosine ramped sine wave tones with mel-spaced center frequencies that matched our sentence spectrograms. These center frequencies ranged from 74.5 Hz to 8kHz. Pure tones were played at 3 intensity levels at 10 dB spacing, with the lowest intensity calibrated to be minimally audible in the hospital room. Each pure tone frequency/intensity pair was repeated 3 times, and inter-stimulus intervals were jittered (range 0.28 s minimum ISI – 0.5 s maximum ISI).

### 4.5 Electrode selection

We identified electrodes with robust responses to speech sounds using a bootstrap t-test. For each electrode, we compared random 1-second chunks of neural data during speech stimuli to 1-second chunks during silence for 10,000 bootstrap iterations (with replacement). With each iteration, we computed the mean difference in activity for speech vs. silence, and then calculated the bootstrapped p-value as the mean number of times this difference in activity was greater than zero. We then selected electrodes with a significant response to speech at a threshold of *p* < 0.001. This resulted in a total of 2,100 speech-responsive electrodes across the 27 patients.

### 4.6 Soft clustering of time series data using convex non-negative matrix factorization

We used convex non-negative matrix factorization (cNMF) (Ding et al., 2010) to uncover functional areas based on correlated activity during a natural speech listening task. In brief, we estimated the time series X [n time points x p electrodes] with the following factorization:

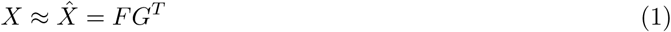

where *F* = *XW*.

The G matrix [*p* electrodes × *k* clusters] represents the spatial weighting of an electrode on a given cluster, and the W matrix [*p* electrodes × *k* clusters] represents the weights on each of the electrode time series. Restricting F to be a convex combination of the electrode time series allows us to compute a time series “centroid” – that is, the weighted time series XW for each cluster k will give us the prototypical time series for that cluster (see (Ding et al., 2010) for proof of this concept, and Fig. 1b for cluster time series).

We concatenated the z-scored time series for all 27 subjects and performed the clustering analysis on all subjects simultaneously to find patterns of activity that were consistent across subjects. We restricted this analysis to the sentences that were heard by all subjects, which included a total of 319 sentences. We initialized the W matrix by first performing an eigenvalue decomposition on the unit-normed covariance matrix *X^T^X*, followed by a varimax rotation and rectification. The G matrix was then initialized according to Ding et al. 2008 (Ding et al., 2008), followed by alternating updates of *W* and *G* as described in Ding et al. 2010 (Ding et al., 2010) until convergence was achieved.

To evaluate the number of clusters, we calculated the percent variance explained when projecting the data onto the computed NMF clusters. For this, we used the following equation:

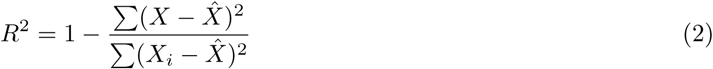

We then plotted the additional percent variance explained for *k* = 2 to *k* = 32 clusters.

### 4.7 Silhouette index

To evaluate cluster separability, we employed the silhouette index *s*(*i*), which describes how well each electrode *i* is matched to its own cluster compared to the non-match cluster. This takes the form:

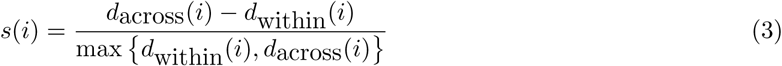

where *d*_across_(*i*) is the lowest average dissimilarity of electrode *i* to the cluster for which it is not a member (as measured by the squared Euclidean distance), and *d*_within_(*i*) is the average dissimilarity of electrode *i* with all other electrodes in the same cluster. The silhouette index was calculated within each subject separately. For the functional clustering, we calculated the dissimilarity as the squared Euclidean distance between the NMF activation weights G for each cluster. For anatomical clustering, the dissimilarity was calculated using the physical pairwise distance between electrodes within or across clusters.

### 4.8 Trajectory analysis

State-space trajectory analyses were performed by projecting data from all electrodes onto NMF basis functions. This is equivalent to the calculation of the cluster time series “centroid” *F*, described above. For Zone 1, this was the weighted time series *XW*_1_, where only the first column of W was used, and for Zone 2, this was the weighted time series *XW*_2_, with only the second column of *W*. For analyses that were restricted by subject, we used only the columns of *X* and rows of *W* corresponding to electrodes within the subject of interest, and calculated *XW* for those electrode subsets.

### 4.9 Spectrotemporal and feature-temporal receptive field estimation

To model acoustic and phonetic transformations in primary and non-primary areas, we used linear encoding models to describe the high gamma activity recorded at each electrode as a weighted sum of stimulus features over time. This model is known in the literature as the spectrotemporal receptive field, and is widely used to describe selectivity for natural stimuli (Theunissen et al., 2001). The models were of the form:

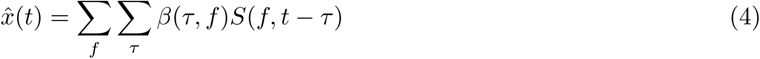

Where *x* is the neural activity recorded at a single electrode, *β*(*τ*, *f*) contains the regression weights for each feature *f* at time lag *τ*, and S is the stimulus representation. We estimated models using two representations of the data: (1) a spectrogram-based representation, and (2) a phoneme feature based representation. For the spectrotemporal stimulus representation, we used the mel-band spectrogram as in our previous work (Mesgarani et al., 2014). The mel band frequencies ranged from approximately 75 Hz to 8 kHz, using an auditory filter bank with a cosine transform that gives a representation of spectral power over time that mimics the filtering performed by the human auditory system (Slaney, 1998).

For the phoneme feature representation, we constructed a binary phoneme feature matrix describing each sentence as a set of features (1 for the presence of a feature, and 0 for its absence) describing phonetic or other linguistic content. Based on previous work showing that the STG responds to phonetic features rather than single phonemes (Mesgarani et al., 2014), we collapsed the phoneme transcription from TIMIT to phonetic features, including voiced and unvoiced fricatives, voiced and unvoiced plosives, liquids/nasals/glides, and vowels. Vowels were split by stress defined at three levels (P1=unstressed, P2=medium stress, P3=high stress). To model overall amplitude envelope fluctuations, we included a feature for speech vs. silence. To model response nonlinearities at the beginning of sentences and after pauses, we also included a sentence onset feature to mark the first phoneme of each sentence, as well as features marking the end of long pauses (> 200 ms) or short pauses (< 200 ms). Finally, the tonic syllable was taken as the final full-stress syllable of the sentence (Halliday, 1963; Ladefoged and Johnson, 2011). The final stressed syllable of a typical English sentence or phrase carries special prosodic significance (Kingdon, 1958; Palmer, 1922; Wells, 2006), for example it begins the terminal contour (Hockett, 1958), including the famous upswing of many questions (Butler, 1634), and constitutes the peak of information delivery (Hultzén, 1959) within a prosodic phrase. This was always within a content word (as opposed to functional words), and only rarely (<≈ 10 sentences) did manual identification of the tonic syllable differ. These were case where, due to sentence stress, an earlier stressed syllable was perhaps equal or greater in prosodic prominence as the final stressed syllable. However, linguists disagree as to whether the sentence-stress syllable should necessarily be considered the tonic syllable, and we note that sentence stress is certainly not an all-or-none phenomenon. Thus, assignment of the tonic syllable by listening for sentence stress will necessarily be highly subjective and time-consuming. Since only a handful of sentences were at question, which did not affect the results, it was deemed better to stick with the simple rule-based tonic syllable (i.e., the last full-stress syllable of the sentence), which can be applied automatically and reproduced easily across laboratories.

We fit receptive fields using time delays of up to 600 ms in order to account for the longer responses observed in the anterior STG. *β* weights were fit using ridge regression, where the ridge parameter was estimated using a bootstrap procedure in which the training set was randomly divided into 80% prediction and 20% ridge testing sets. The ridge parameter was chosen as the parameter that gave the best average performance across electrodes as assessed by correlation between the predicted and ridge test set performance. The final performance of the model was computed on a final held out set not included in the ridge parameter selection. Performance was measured as the correlation between the predicted response on the model and the actual high gamma measured for sentences in the test set.

### 4.10 Modulation transfer function analysis

We calculated the modulation transfer function (MTF) of each STRF as the 2D Fourier transform of the STRF (Singh and Theunissen, 2003). After taking the 2D Fourier transform, values were squared, log transformed, and multiplied by 10 to convert units to power (dB). The best temporal modulation and best spectral modulation were then jointly selected by identifying the peak of the MTF matrix, as described previously (Hullett et al., 2016).

### 4.11 Response latency analysis

To calculate the response latencies from the STRF, we calculated a temporal kernel by taking the mean across all frequencies in the STRF matrix. We calculated the onset latency as the time at which the derivative of the temporal kernel reached its maximum. The peak latency was the peak of this temporal kernel, and the offset latency was the time after the peak at which the derivative of the temporal kernel was maximally negative.

### 4.12 Statistical tests

For data that deviated from normality, we used nonparametric Wilcoxon rank sum (for unpaired data) or signed rank tests (for paired data). In some cases, a bootstrap t-test was used.

## 5 Acknowledgments

The authors would like to thank Dr. Michael Lawton for assistance with data collection in intraoperative recordings. The authors would like to thank Christoph Schreiner, Keith Johnson, Michael Stryker, and members of the Chang Lab for helpful comments on the manuscript. This work was supported by grants from the NIH (F32 DC014192-01 Ruth L. Kirschstein postdoctoral fellowship, to LSH, and DP2-0D00862 and R01-DC012379 to EFC). E.F.C is a New York Stem Cell Foundation-Robertson Investigator. This research was also supported by The New York Stem Cell Foundation, The McKnight Foundation, The Shurl and Kay Curci Foundation, and The William K. Bowes Foundation. We gratefully acknowledge the support of NVIDIA Corporation with the donation of the Tesla K40 GPU used for this research.

## 6 Author Contributions

LSH, EE, and EFC conceived of and designed the experiment. LSH, EFC, and others collected the data. LSH and EE analyzed the data. EC performed surgery and grid implantation. LSH, EE, and EFC wrote the paper. The authors report no conflict of interest.

## 8 Supplementary Information

### 8.1 Supplementary Methods

*Sentence control stimuli*. In addition to natural sentences and pure tones, a subset of subjects were presented with a set of control stimuli that were synthesized from 10 of the original TIMIT sentences. These 10 sentences were the subset that were repeated 10-20 times, and included 5 male and 5 female speakers. Control conditions included time-reversed sentences and spectrally rotated sentences. Time-reversed sentences were constructed by flipping the stimulus waveforms such that each sentence was played backwards in time from its original version. Spectrally rotated sentences were constructed according to methods described by (Blesser, 1972).

### 8.2 Supplementary Tables and Figures

**Table 1:** Clinical and demographic details for subjects. Hem = hemisphere of implantation.

| Subject | Hem | Age | Sex | Handedness | Language dominance | Epilepsy focus |
| --- | --- | --- | --- | --- | --- | --- |
| EC36 | L | 32 | F | R | L | Left anterior temporal lobe |
| EC37 | L | 25 | F | L | L | Left medial temporal lobe |
| EC55 | L | 31 | F | R | L | Left hippocampus/anterior lateral temporal |
| EC60 | L | 20 | F | R | L | Left hippocampus |
| EC63 | L | 41 | F | R | L | Left anterior temporal lobe |
| EC2 | L | 44 | M | R | L | Left posterior STG |
| EC33 | L | 19 | M | R | L | Left anterior frontal lobe |
| EC35 | L |  | M |  | L | Left anterior temporal lobe |
| EC6 | R | 29 | F | L | R | Right temporal mesial lobe |
| EC52 | R | 53 | F | R | L | Right anterior/superior frontal lobe |
| EC61 | R | 47 | F | R | L | Right anterior mesial basal temporal lobe/anterior lateral temporal |
| EC77 | R | 44 | F | R | L | Right amygdala/inferior temporal cortex |
| EC79 | R | 47 | F | R | L | Right mesio-temporal, middle temporal, middle-lateral temporal gyrus |
| EC82 | L | 60 | M | R | L | Left mesial temporal structures |
| EC84 | R | 30 | M | R | L | Right anterior temporal lobe |
| EC96 | R | 32 | F | L | L | Right anterior medial temporal |
| EC92 | L | 26 | M | R | L | Left mesial and anterior lateral temporal cortex |
| EC113 | L | 22 | M | R | L | Left anterior temporal lobe |
| EC118 | L | 31 | F | R | L | Left hippocampus/amygdala |
| EC129 | R | 33 | F | R | L | Posterior aspect of the right superior frontal gyrus |
| EC131 | L | 28 | M | R | L | Left anterior and mesial temporal regions |
| EC28 | R |  | M | R | L | Left hippocampus |
| EC56 | R | 56 | M | R | L | Right temporal and temporal frontal junction |
| EC53 | L | 52 | M | R | L | Left insular tumor |
| EC75 | L | 29 | M | R | L | Left frontal operculum tumor |
| EC85 | L | 26 | M | R | L | Left frontal operculum tumor |
| EC89 | L | 43 | M | R | L | Left insular tumor |

**Figure Supplement 1:**
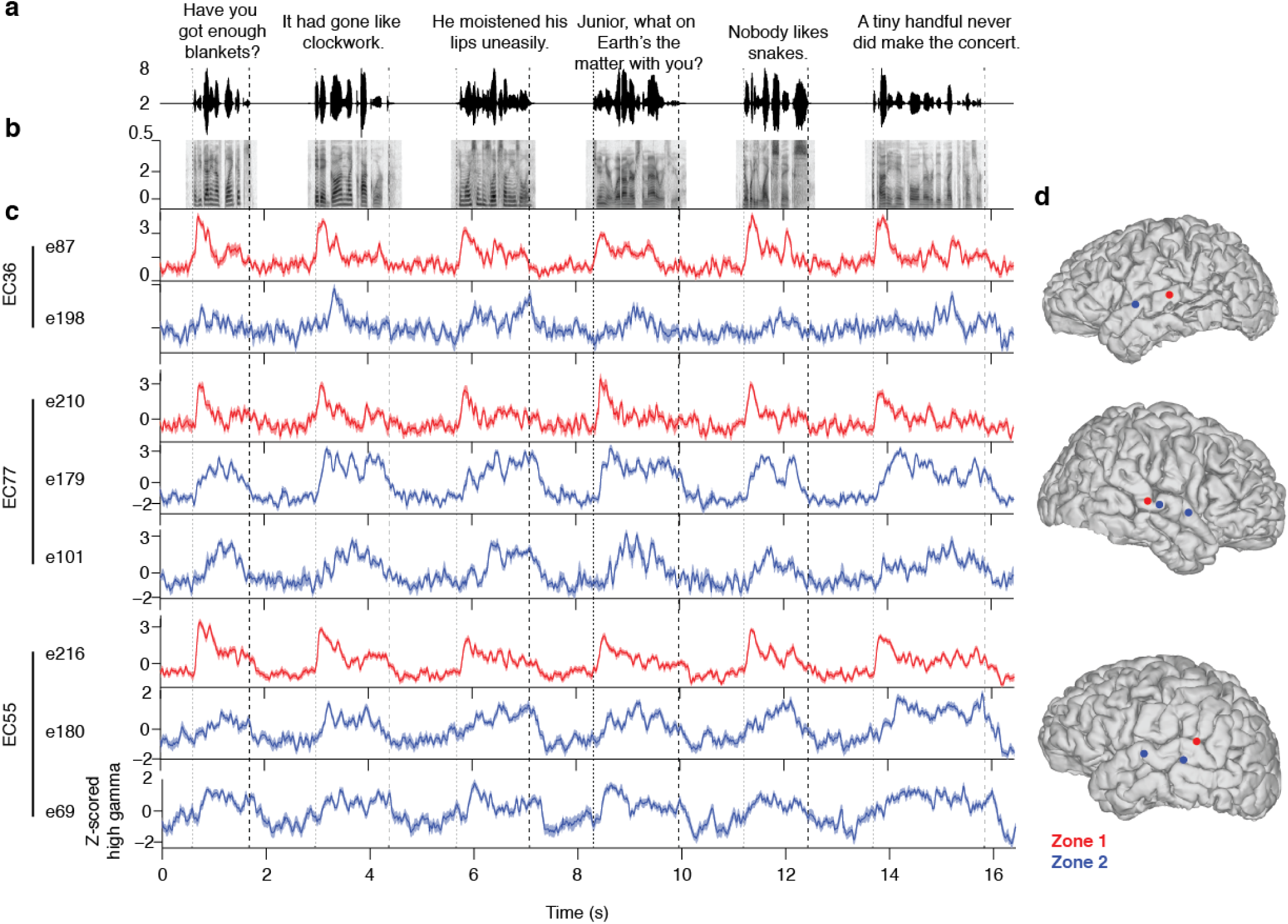
Single channel examples of responses within Zone 1 (red) and Zone 2 (blue) electrodes in 3 example subjects. Zone 1 electrodes show strong responses to sentence onsets, whereas Zone 2 electrodes show a more sustained response throughout the sentence. (a) Waveforms for six sentences from the TIMIT database. The onset of the first phoneme and the offset of the last phoneme in each sentence are marked as dashed lines. Note that these sentences contain very different phonetic content, especially at sentence onset. (b) Mel-band spectrograms of same sentences in (a). (c) Example Z-scored high gamma average responses for 8 electrodes from 3 subjects, classified as belonging to the Zone 1 (red) and Zone 2 (blue) group. Note that in all three subjects, the Zone 1 electrodes respond strongly at sentence onset, regardless of the starting phoneme. Zone 2 electrodes, on the other hand, do not show a strong onset response and show a more even response throughout the sentence. Electrode traces for each subject are organized from posterior to anterior. (d) Anatomical locations of electrodes shown in (c).

**Figure Supplement 2:**
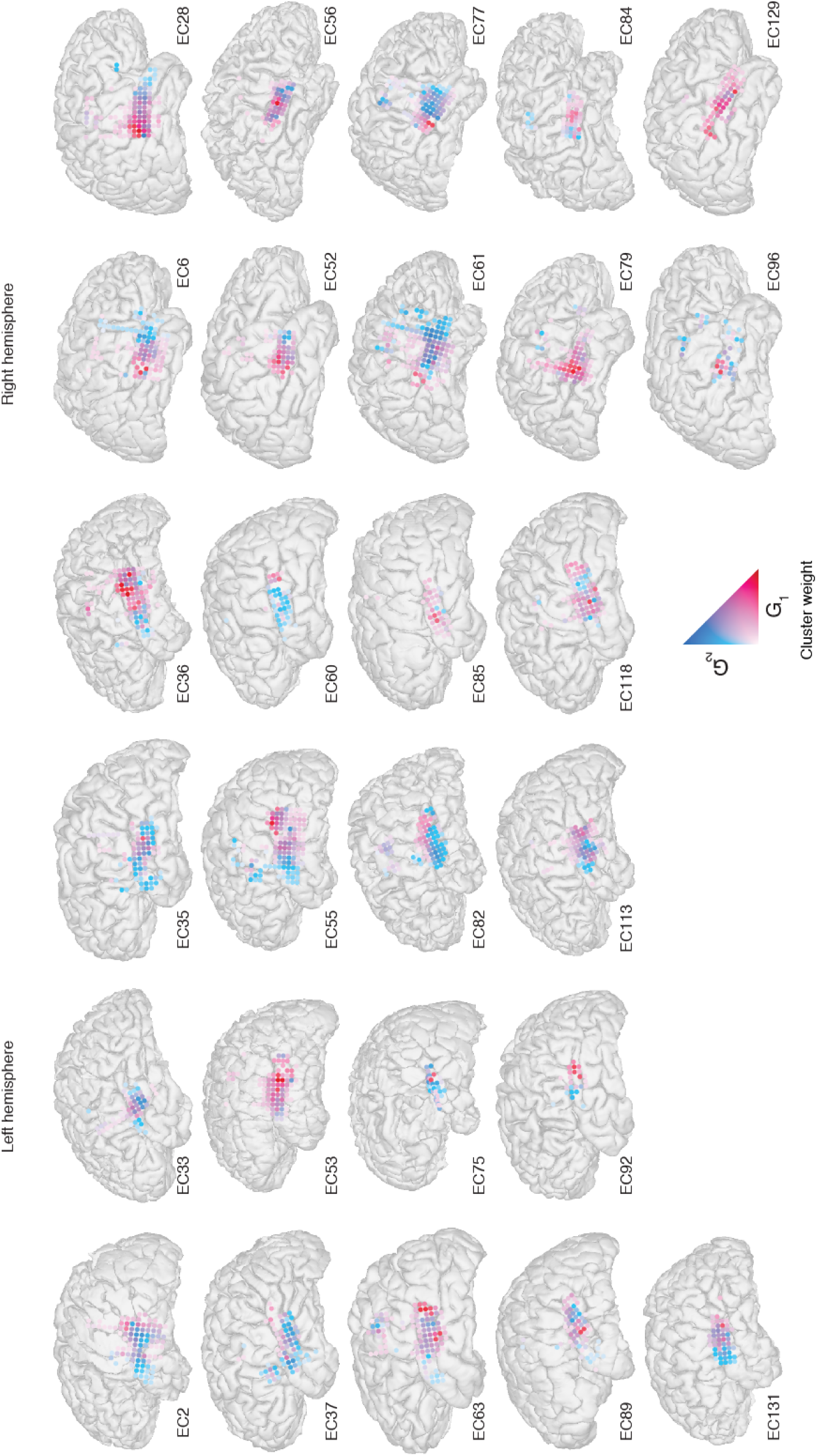
NMF clustering results on all subjects included in this study. Speech responsive electrodes are shown and colored according to the weight on NMF cluster 1 (Zone 1, red) or NMF cluster 2 (Zone 2, blue).

**Figure Supplement 3:**
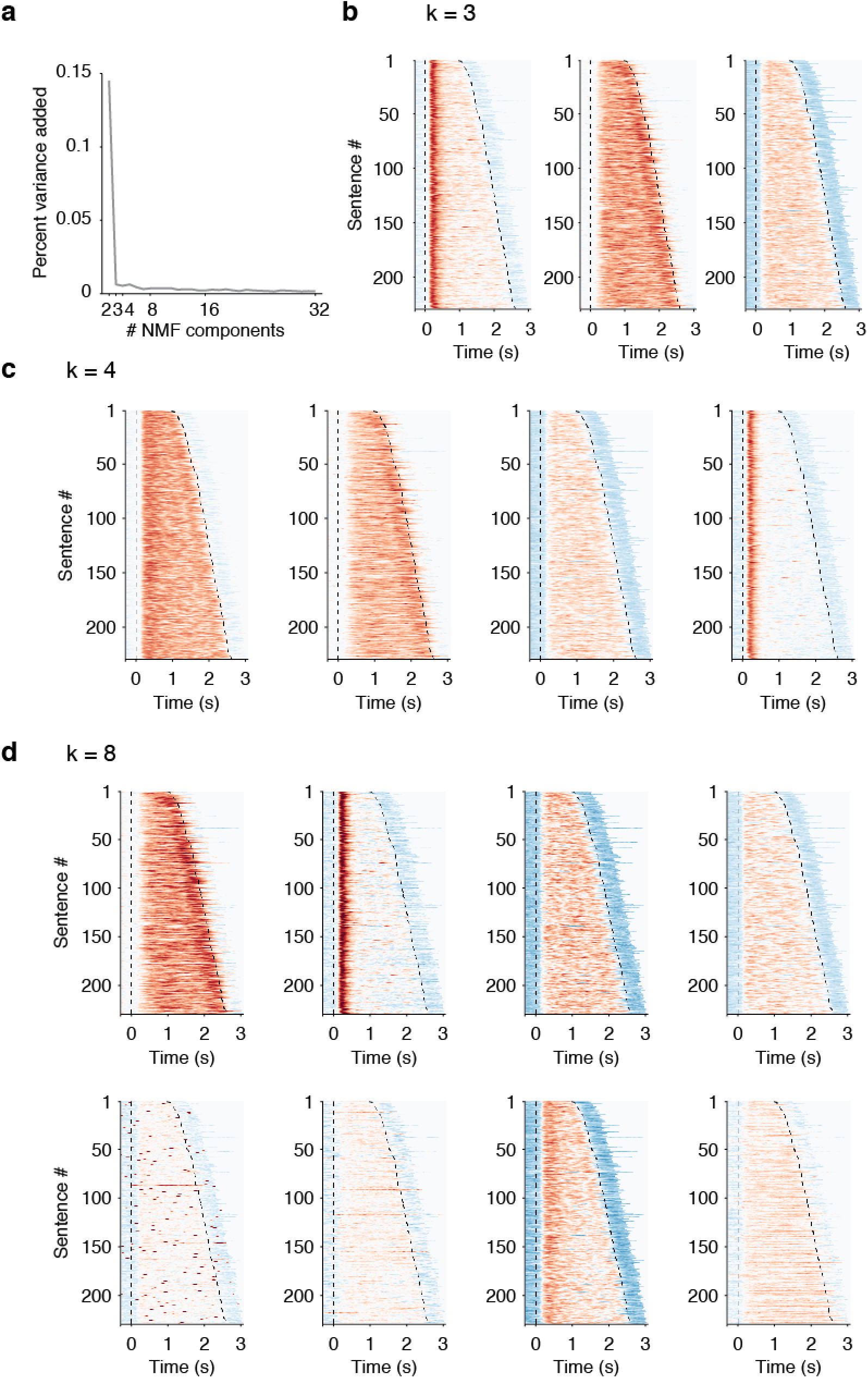
NMF clustering results for additional numbers of clusters across all patients. (a) Adding clusters beyond k=2 only marginally improves reconstructions (as measured by percent variance explained across all data). Percent variance added as calculated as the difference between the percent variance explained for each successive number of clusters. (b) Clustering results for k=3 clusters. (b) Results for k=4 clusters. (c) Results for k=8 clusters. In all cases, variants of the “Zone 1” and “Zone 2”-type responses were observed, with varying degrees of responsivity.

**Figure Supplement 4:**
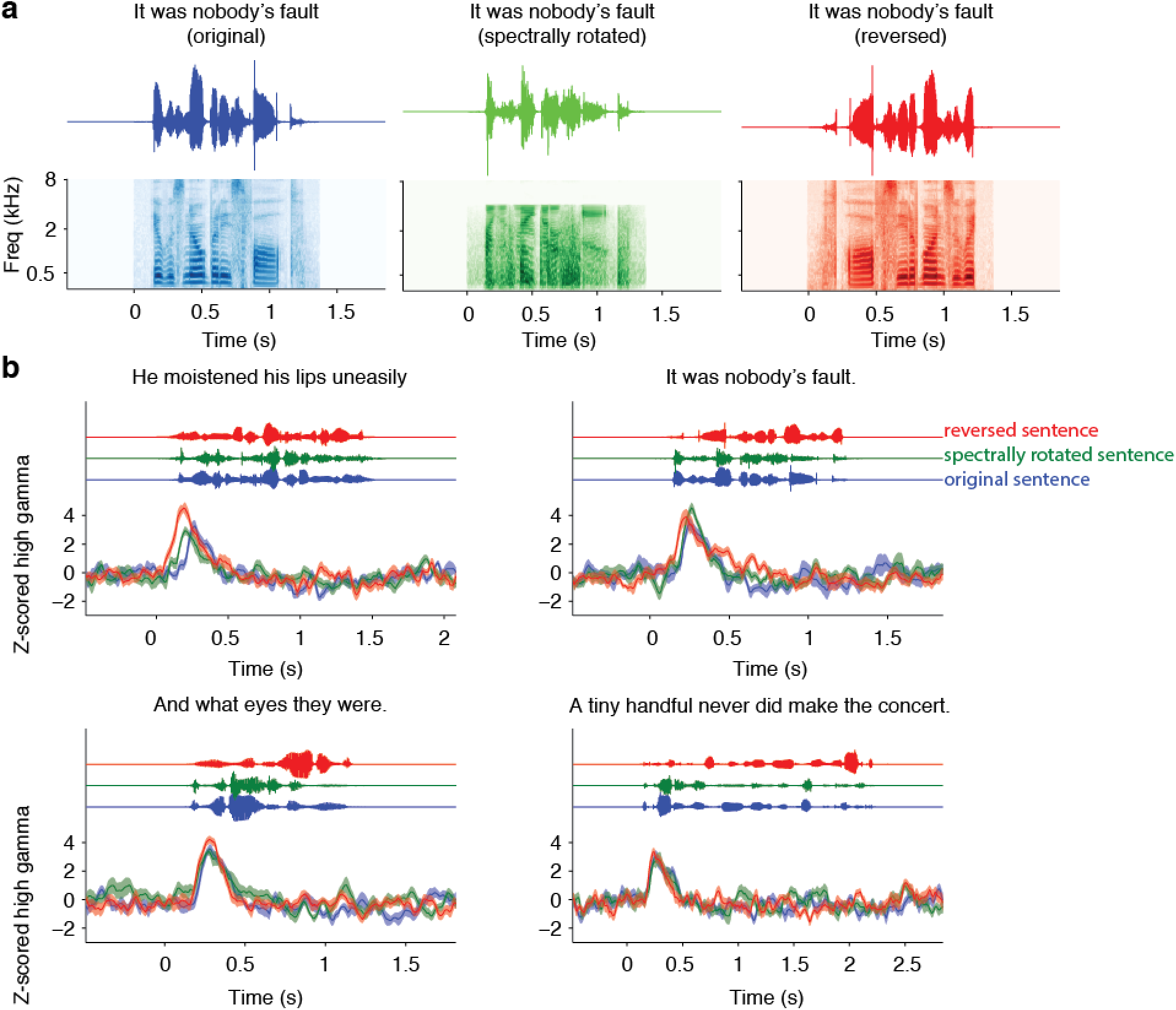
(a) Example waveform and spectrograms for spectrally rotated and reversed control stimuli synthesized from the original sentence at left (“It was nobody’s fault.”) (b) Onset responses for a Zone 1 electrode in response to original, spectrally rotated, and reversed sentences. Original sentence text is specified at the top of each subplot, along with the stimulus waveform at the top of the subplot, and Z-scored high gamma activity in response to these stimuli across 10 repetitions of each sentence. We observed strong onset responses in all stimulus types, suggesting that these onset responses are not speech-specific.

**Figure Supplement 5:**
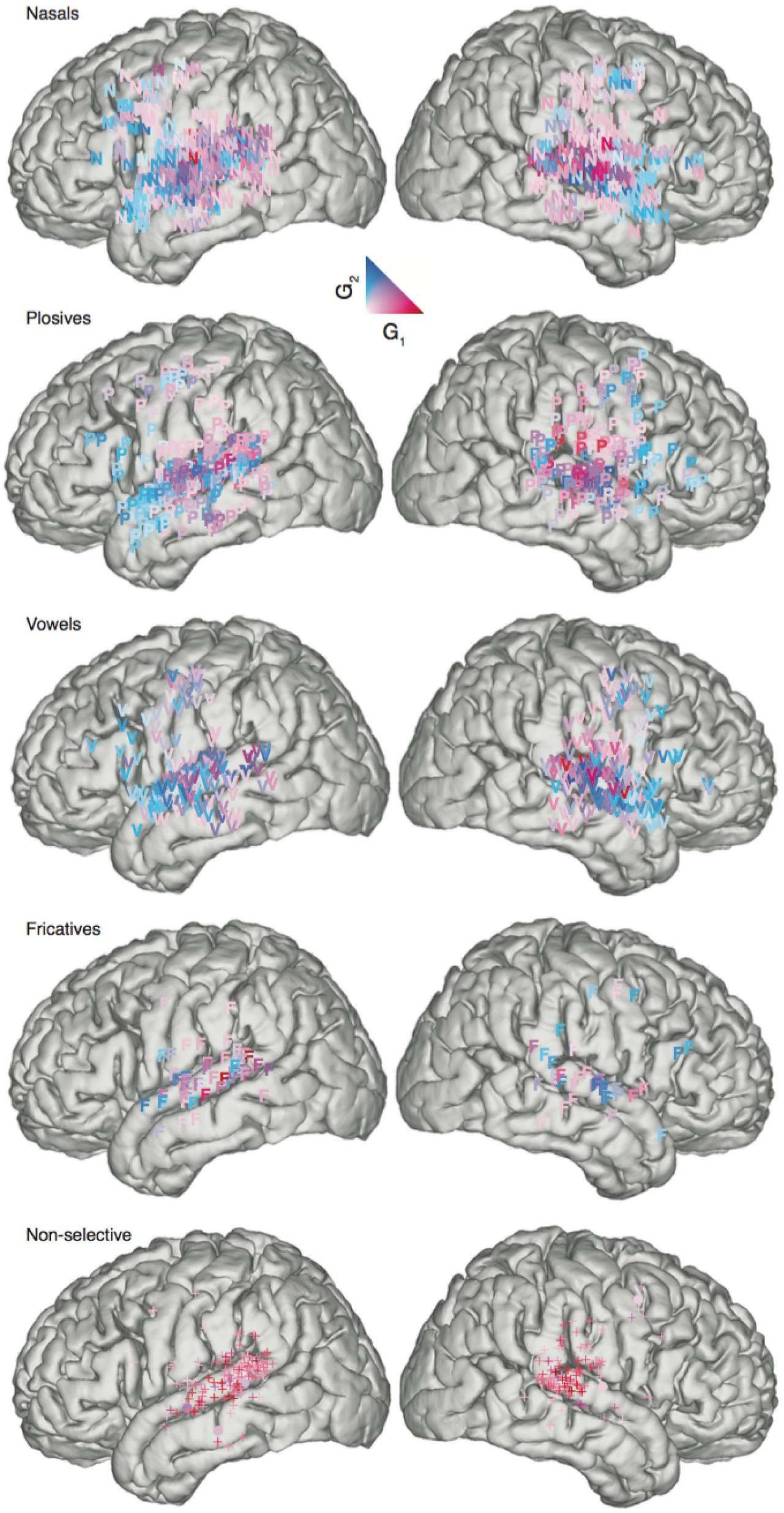
No evidence for clustering of phoneme features within Zone 1 and Zone 2. Phoneme feature maps across all subjects, colored by weight on Zone 1 (red) vs. Zone 2 (blue). N = nasals, P = plosives, V = vowels, F = fricative, + or • = non-selective Zone 1 or Zone 2. Only non-selective (mostly onset) responses were localized to a particular area of speech cortex – most of these responses were in posterior STG.

**Figure Supplement 6:**
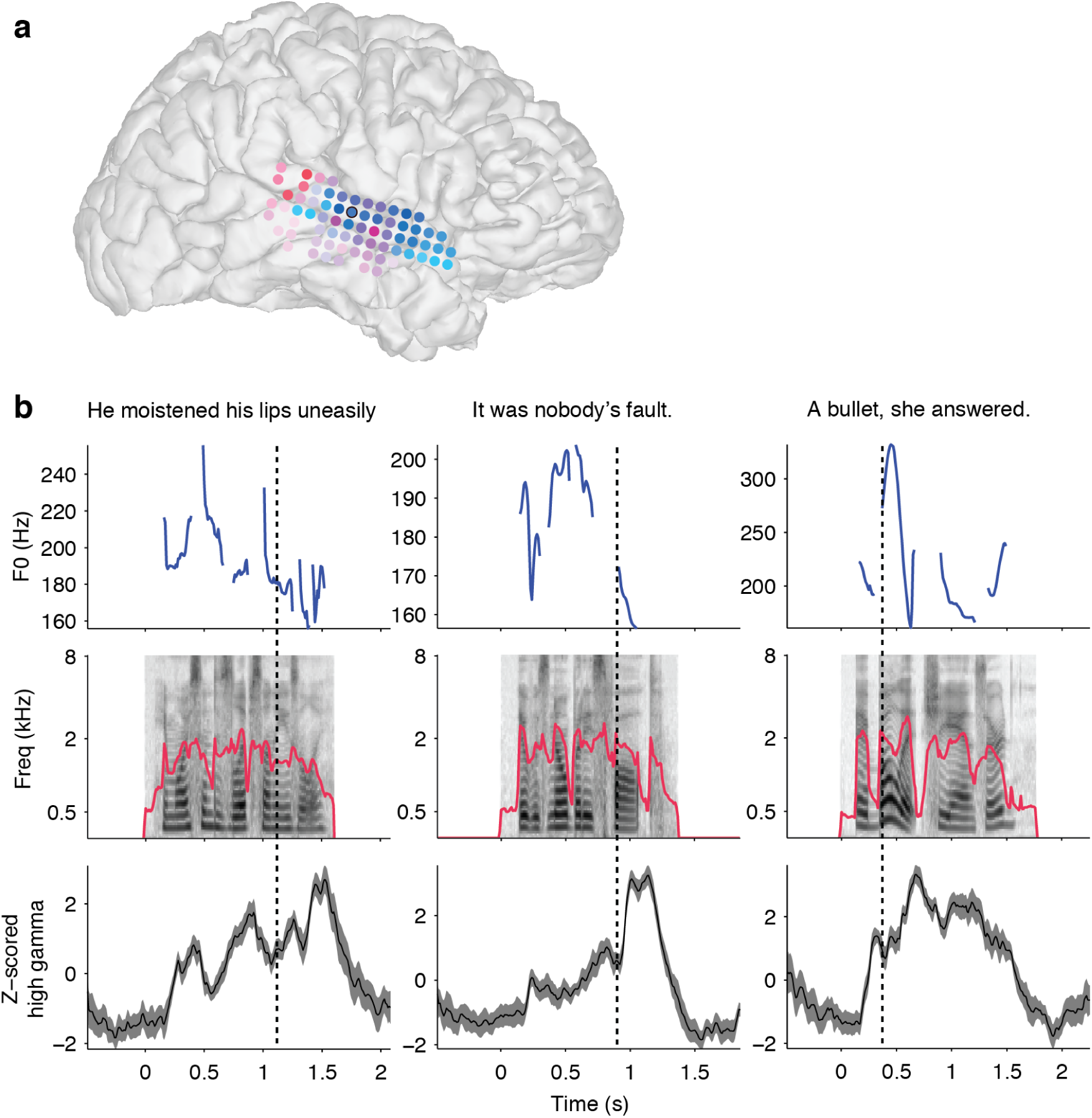
Example responses of a tonic-syllable selective Zone 2 electrode to TIMIT sentences. (a) Reconstruction of subject’s brain, with weight on Zone 1 colored red and weight on Zone 2 colored blue. The electrode whose activity is shown in (b) is outlined in black. (b) Response of electrode in (a) to three example sentences, with tonic syllable marked as a dashed line.

**Figure Supplement 7:**
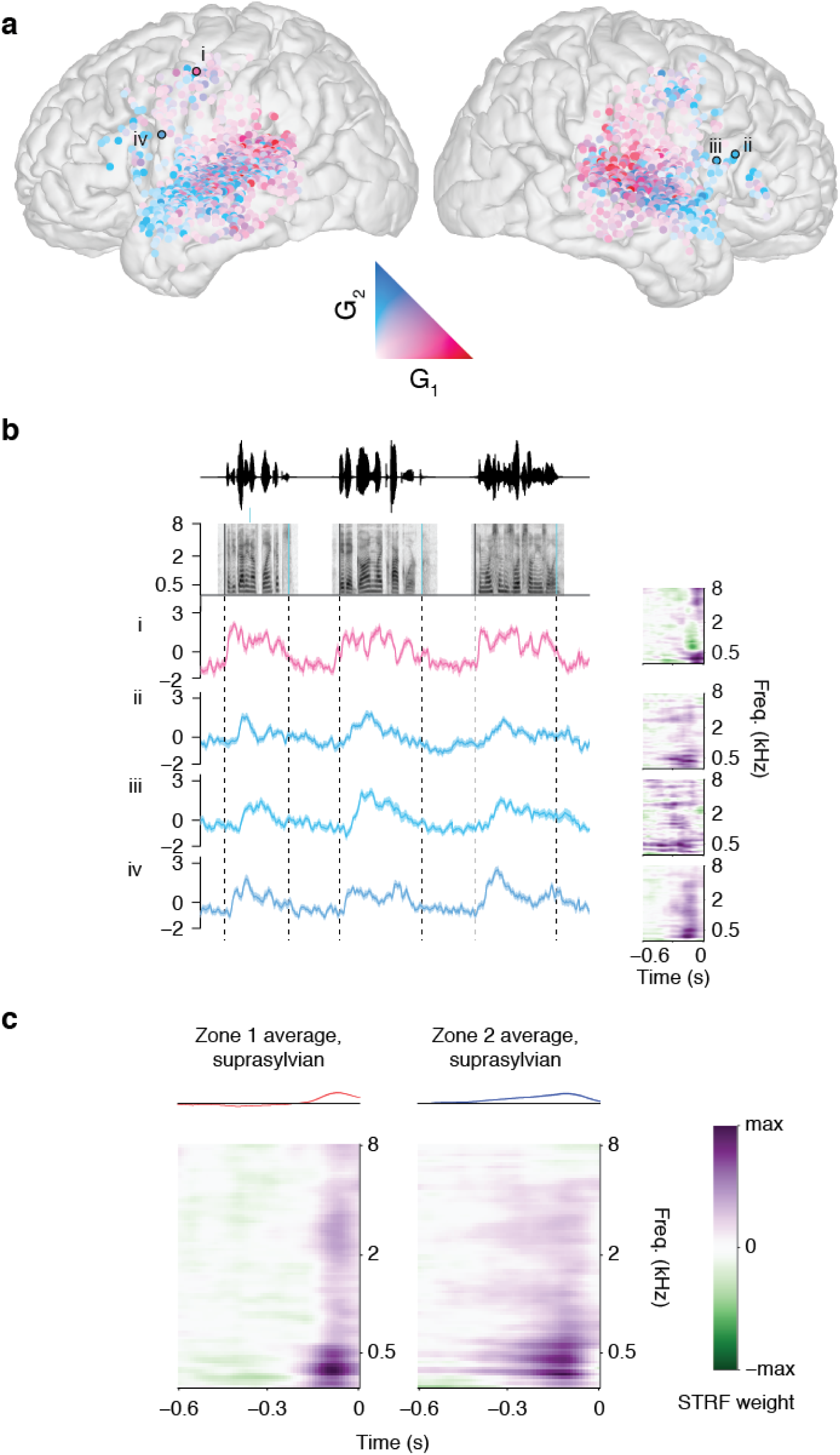
Zone 1 and Zone 2 response types are an organizing feature of suprasylvian areas of the speech-responsive cortex. (a) Electrodes on the STG and in suprasylvian areas, colored by their weight on Zone 1 (red) or Zone 2 (blue). (b) Example single electrode responses and their STRFs (right) for suprasylvian electrodes in Zone 1 (red/pink) and Zone 2 (blue). Zone 1 electrodes showed similar strong onset responses, whereas Zone 2 electrodes responded to speech features throughout the sentence rather than primarily at onset. (c) Average Zone 1 and Zone 2 STRFs show a temporal profile similar to Zone 1 and Zone 2 electrodes in STG (Fig. 2a) and on the temporal plane (Fig. 5f).

**Figure Supplement 8:**
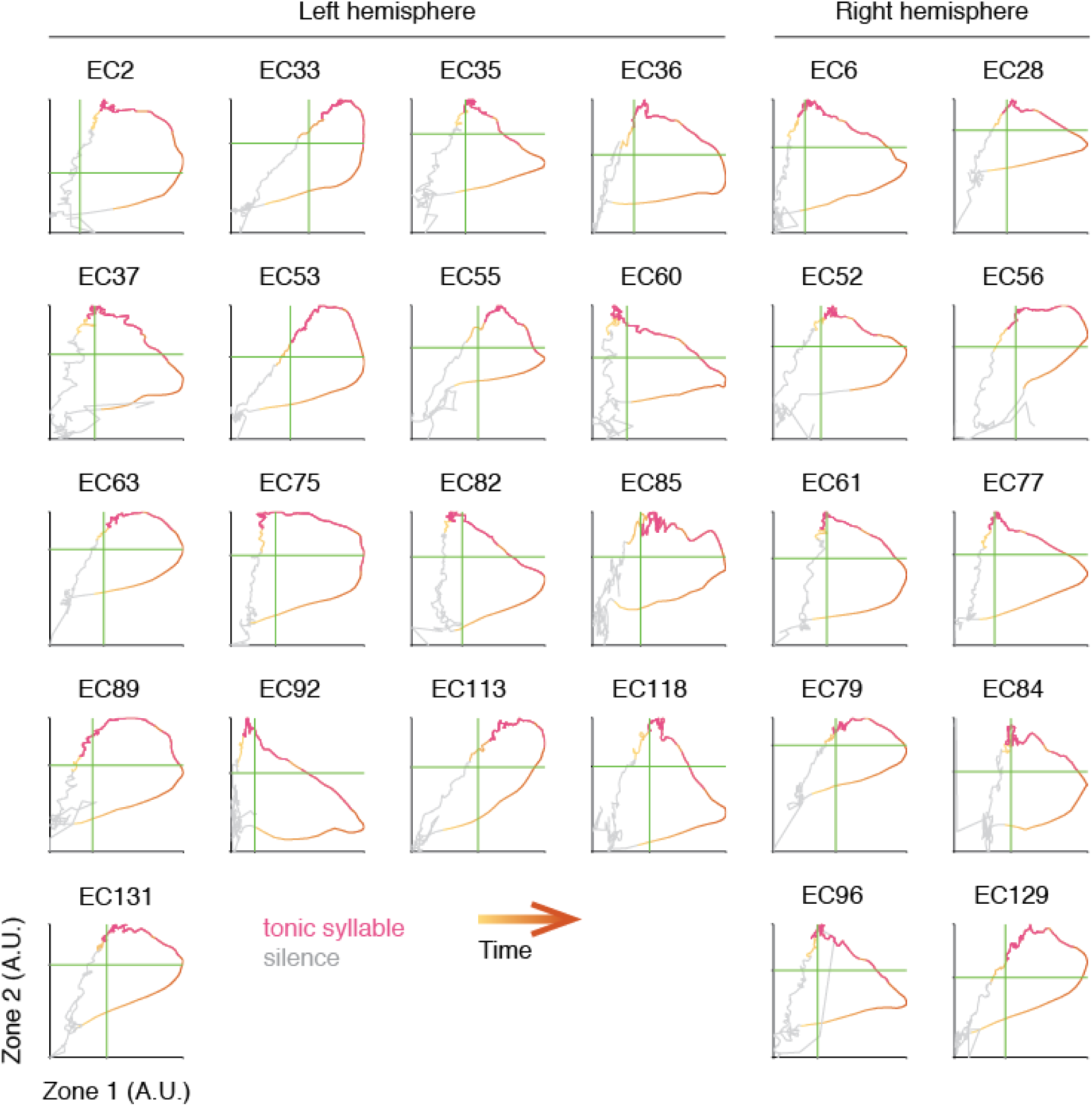
State space trajectories within single subjects. For each subject, the time series is projected onto Zone 1 and Zone 2 electrodes and shown averaged across all sentences. Trace changes color from yellow to orange over time during the sentence. Gray indicates silent portions of the stimulus. All possible positions of the tonic syllable are plotted in pink.

